# Time-resolved single-cell screening reveals the dynamic regulatory architecture of human cardiogenesis

**DOI:** 10.64898/2026.11.03.764059

**Authors:** D. Lindenhofer, K. Mikulik, S. Mantz, H. Jung, K. Ueltzhöffer, M. Kueblbeck, L. Rueda-Gensini, A. Muñoz-Verdú, J. Haas, B. Meder, D. Schraivogel, M. Mall, W. Huber, A. Pettet, J. P. Reddington, B. Clarke, O. Stegle, L. M. Steinmetz

## Abstract

Congenital heart disease (CHD) originates during early embryogenesis, yet when disease genes act cannot be determined from existing data. Human fetal heart atlases lack critical time points, and perturbation screens that systematically assess gene function currently profile at most a few time points. Here we combined single-cell CRISPR screening of 400 disease-relevant genes in cardioids, sampled daily over seven developmental days (∼975,000 cells), with a matched multiome atlas. Perturbation phenotypes were strongly dependent on developmental stage. Perturbations that reshaped cell-fate allocation or gene expression were enriched for CHD-linked genes, and most acted already during progenitor stages. Timing was largely independent of the target gene’s own expression, marking instead when a cell state became functionally dependent on it, and some effects persisted after expression declined. Consensus gene regulatory networks integrating perturbation and multiome data recovered stage-specific regulators and nominated *HOXB2* and *HOXB3* as candidate cardiac regulators. Many perturbations converged on shared downstream programmes, which were reorganised through differentiation. Effects correlated in progenitors were maintained in differentiated cells, split into distinct programmes, converged with other perturbations or disappeared entirely, indicating that gene function is highly context dependent. Beyond advancing our understanding of the developmental regulatory logic underlying CHD, our work provides a resource for developing context-dependent models for *in silico* prediction of perturbation effects.

## Main

Development is a dynamic process in which pluripotent progenitors are specified into differentiated cell types through intermediate progenitors^1^. This process is driven by signalling cues and chromatin changes manifested in cell type-and cell state-specific gene regulatory networks (GRNs)^2,3^. In the developing heart, understanding these transitions remains challenging because existing human fetal heart atlases do not fully capture the critical early window during which specified mesoderm gives rise to distinct cardiac cell types through intermediate progenitors^4–14^. Single-cell perturbation studies can identify regulators of GRNs and link their effects to specific cell types throughout differentiation and to disease-associated genes^6,9,14–18^. It remains unclear whether gene disruption has fixed effects or whether these are redefined by the distinct molecular context of each differentiation stage. Existing single-cell screens assay a limited set of perturbations or time points, thus offering only a partial view of gene function and providing limited insights into how context-specific perturbation effects evolve across differentiation^19–22^.

Self-organising human cardioids provide experimental access to the lineage-defining stages of heart development missing from human heart atlases, modelling mesoderm specification from 18 days post fertilisation (dpf) onwards^23–25^. They generate differentiated cell types within eight days of induction, which renders them an attractive model to study the time-resolved effects of perturbations through temporal sampling and pooled CRISPR/Cas9 screening^20,24–31^.

Congenital heart disease (CHD) is the most common human birth defect, affecting ∼1% of newborns^26^. Diverse cell types have been implicated in CHD pathogenesis by current single-cell fetal heart atlases, including cardiomyocytes, cardiac fibroblasts and endothelial cells^27^. Because CHD arises during early embryogenesis, cardioids have the capacity to uncover CHD origins through understanding the early stages of heart development and associated cell-type-specific regulatory programmes^13,26,27,32–37^. In addition, adult-onset cardiomyopathies often share these developmental origins, as many disease genes act both during cardiac development and in maintaining adult heart physiology.

To resolve the order of events through which CHD and cardiomyopathy genes shape early heart development, we performed a pooled single-cell CRISPR screen of 400 genes in human cardioids, including disease-associated genes and cell-type-specific transcription factors (TFs), sampling daily from cardiac mesoderm to differentiated cell types. To place these perturbation effects in their chromatin accessibility-driven regulatory context, we also generated a matched single-nucleus multiome atlas of unperturbed cardioids.

Our analysis revealed that many disease-associated perturbations already affected the germ-layer and early progenitor stages, exerting molecular consequences prior to the establishment of cardiac lineages. We investigated the timing and regulatory interaction of perturbations, uncovering a hierarchy of both converging and diversifying effects across cell differentiation. Finally, we mapped the regulators of cardiac cell types by integrating perturbational data with single-nucleus multiomics, nominating *HOXB2* and *HOXB3* as candidate cardiac regulators. Collectively our findings reveal an underappreciated role of timing of regulation in CHD, as well as fine-granular relationships between disease genes and pathways.

## Results

### A time-resolved perturbation atlas of human cardioid development

We generated a time-resolved resource of cardioid development consisting of (i) a reference atlas using scRNA-seq and snATAC/RNA-seq (snMultiome-seq), and (ii) a temporally matched pooled CRISPR/Cas9 scRNA-seq dataset (scPerturb-seq)^38^ targeting 400 disease-associated genes and cell-state regulators (Fig. 1a, Supplementary Table 1). Cardioids were generated from WTC-11 induced pluripotent stem cells (iPSCs) using defined patterning cues and densely sampled throughout differentiation from cardiac mesoderm to major cardiac cell types (Fig. 1a, Extended Data Fig. 1a-c)^25^. In total, four independent differentiations were performed: two for the time-resolved scPerturb-seq and one each for the scRNA-seq and snMultiome-seq reference atlas. Following quality filtering, we obtained 22,278 scRNA-seq, 40,088 snMultiome-seq and 974,851 gRNA-assigned scPerturb-seq cells (Fig. 1a, Extended Data Fig. 1d; Supplementary Tables 2 and 3). Cells were annotated using established marker genes^24,25,27,39–42^, and pseudotime was inferred with Palantir^43^ (Fig. 1b-d, Extended Data Fig. 1e, f; Supplementary Table 4).

**Figure 1.**
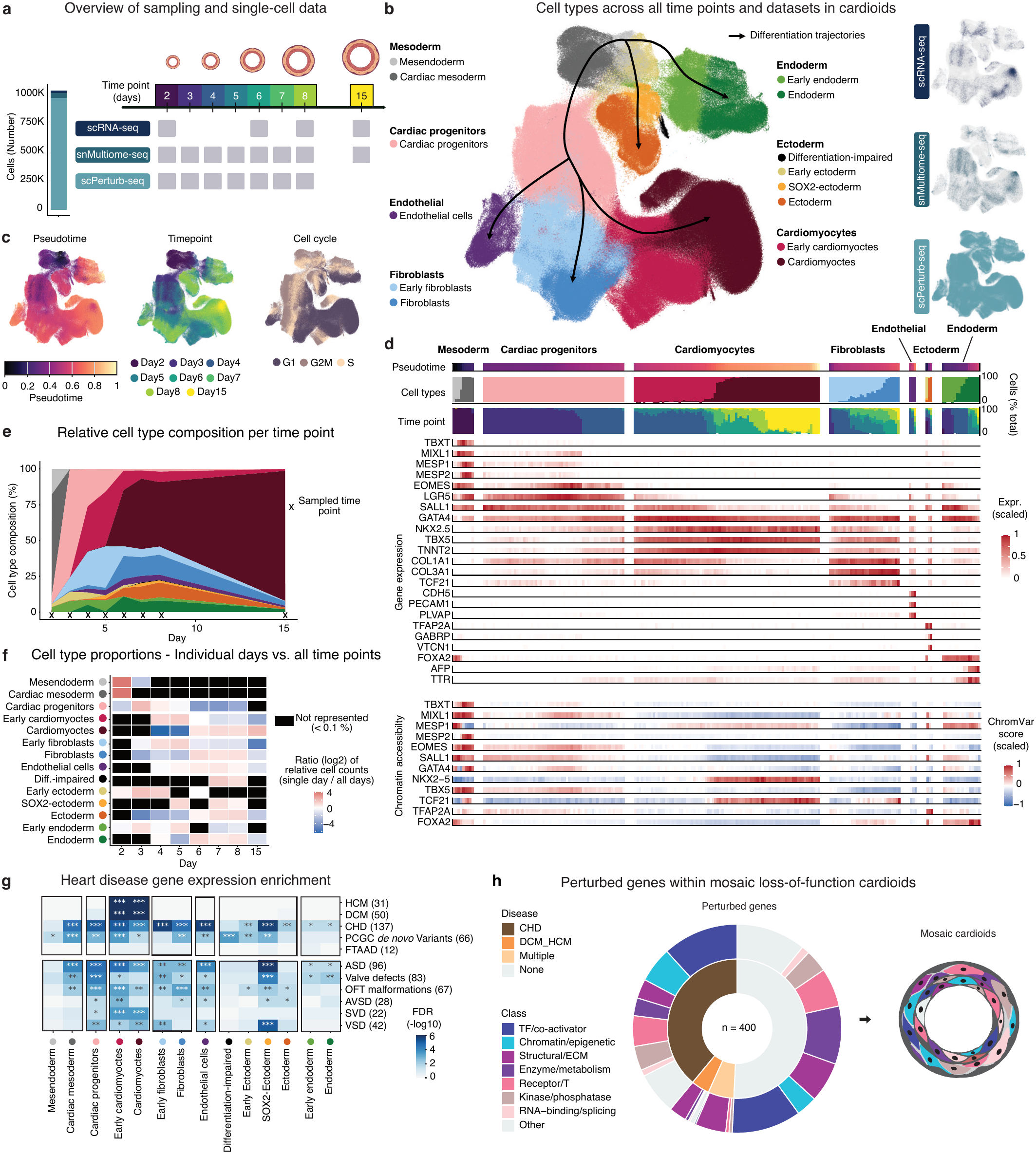
A time-resolved single-cell perturbation dataset in cardioids. a, Experimental design. iPSC-derived cardioids were sampled at the indicated time points for unperturbed reference datasets (scRNA-seq, snMultiome-seq) and for single-cell CRISPR perturbation dataset (scPerturb-seq) of mosaic loss-of-function cardioids (400 target genes)**. b,** UMAP embedding of cells from all three integrated datasets, coloured by harmonized cell-type annotation; inset, contribution of each dataset. Arrows indicate branching differentiation trajectories. **c,** UMAP embedding of cells from all three integrated datasets, coloured by palantir pseudotime, sampling time points and predicted cell-cycle phase. **d,** Pseudotime with sampling time points, cell type, gene expression and chromatin accessibility of cells, grouped into pseudotime bins. Each bin shows cell-type and sampling-time-point composition (%), scaled marker-gene expression and chromVAR TF motif accessibility scores. **e,** Stacked area plot of relative proportions of cell types per time point. Sampled time points are indicated with “X”. **f,** Relative abundances (log2 ratio) per cell type of individual time points compared to the total across all time points sampled. Cell types not represented at a given time point (< 0.1% of cells) are shown in black. **g,** Disease gene enrichment among cell-type-preferential genes (Methods). Shown is the enrichment (one-sided Fisher’s exact test) of genes associated with heart traits (DCM, HCM, CHD, and CHD genes carrying *de novo* coding variants from the PCGC) and a vascular trait (FTAAD, upper panel), or with six CHD subtypes (lower panel), among the top 300 most highly and specifically expressed genes of each cell type (specificity z-score > 1, ranked by mean expression). Colour corresponds to Benjamini-Hochberg-adjusted FDR (-log10). *FDR < 0.05, **FDR < 0.01, ***FDR < 0.001. **h,** Overview of the 400 perturbed genes. Inner circle: disease-associated genes. Outer circle: manually annotated functional classes.

Cardioid differentiation involved rapid fate acquisition between days 2 and 6, with early cell types (mesendoderm, cardiac mesoderm, cardiac progenitors) present only transiently during narrow developmental windows from day 2 to day 4 (Fig. 1e). This underscores the value of dense temporal sampling, as sampling at individual time points would have led to missing or drastically over-or underrepresented cell types (Fig. 1f). From day 4, cells expressing cardiomyocyte, fibroblast and endothelial markers emerged. Cardiomyocytes expressed first-heart-field genes, consistent with the patterning cues used^24,25^ (Extended Data Fig. 1e, f). Fibroblasts lacked expression of canonical epicardial markers, indicating non-epicardial origin^39,40^. From day 2 onward, a subset of cells diverged towards endoderm, while a small population acquired a non-neuronal amniotic ectoderm fate from day 2 (*VTCN1, GABRP* and *TFAP2A*) (Extended Data Fig. 1e). A few perturbations yielded differentiation-impaired cells retaining pluripotency gene expression (Extended Data Fig. 1e). Fate probabilities assigned with CellRank2^44^ supported early divergence into mesoderm, endoderm and ectoderm at day 2 (Extended Data Fig. 2a, b). Cardiomyocytes, fibroblasts and endothelial cells were specified through intermediate cardiac progenitors, with prolonged cardiomyocyte-fibroblast plasticity.

Cardioids model early developmental stages and cell types that are largely inaccessible in *in vivo* atlases^27^. To determine when and in which cell types disease genes are expressed, we tested each cell type’s preferentially and highly expressed genes for enrichment of genes linked to hypertrophic cardiomyopathy (HCM), dilated cardiomyopathy (DCM) or CHD, including those with *de novo* coding variants in the Pediatric Cardiac Genomics Consortium cohort (PCGC *de novo*) (Fig. 1g, Supplementary Table 5)^27,36,41^. CHD genes, including genes associated with valve defects, atrial septal defects (ASD) and outflow tract (OFT) malformations, were broadly enriched, including in early cell types, such as cardiac mesoderm and cardiac progenitors, that are restricted to a narrow time window. In contrast, DCM and HCM genes were specifically enriched in cardiomyocytes, consistent with their reported cardiomyocyte-restricted biology^27^. Familial thoracic aortic aneurysm and aortic dissection (FTAAD) genes, a smooth-muscle negative control, were not enriched in any cardiac cell type. Cardioids thus model disease-relevant expression patterns across diverse cell types and disease phenotypes.

The 400 genes for scPerturb-seq were selected based on associations with CHD, cardiomyopathies and brain-heart multi-organ syndromes, prioritising genes expressed in the reference atlas and including five to ten TFs per major cell type (Fig. 1h, Methods, Supplementary Table 6). Genes were disrupted by introducing loss-of-function mutations in iPSCs, thereby perturbing gene function already before the onset of cardioid differentiation. Cas9-expressing iPSCs were transduced with a lentiviral gRNA library (4 gRNAs/gene), including two types of negative control (NC) gRNAs: (i) 16 non-targeting control gRNAs and (ii) 16 gRNAs targeting the CLYBL locus (Extended Data Fig. 2c, Supplementary Tables 1 and 7). To capture perturbation effects across transient cell states from mesoderm induction to differentiated cell types, cardioids were sampled at seven consecutive time points (day 2–8, 24 h apart). After quality control, cells were assigned a unique gRNA (read-count purity ≥90%, ≥3 UMIs, ≥100 reads; Extended Data Fig. 2d-g). With this dataset in hand, we next turned towards the question of linking disease genes to developmental mechanisms in specific cell states and their temporal patterns through differentiation.

### CHD genes impact early cell fate decisions

First, we tested which perturbations shift cell fate by comparing cell-type abundance between perturbed and control cells (Cochran-Mantel-Haenszel test). This identified 41 perturbations that significantly altered cell type proportions with pronounced effects (FDR < 10LL, absolute log2 odds ratio ≥ 0.5 in at least two cell types; Fig. 2a, b). These 41 pronounced cross-cell-type perturbations were significantly enriched for transcriptional/chromatin regulators and CHD-linked genes (Extended Data Fig. 3a, b). To group perturbations with similar effects on cell fate, we clustered them by their effect-size profiles across cell types (log2 odds ratios; hierarchical clustering; Methods), which yielded nine cell-fate clusters (A-I) spanning early germ-layer specification and subsequent cardiac differentiation (Fig. 2b, Extended Data Fig. 3c, Supplementary Table 8).

**Figure 2.**
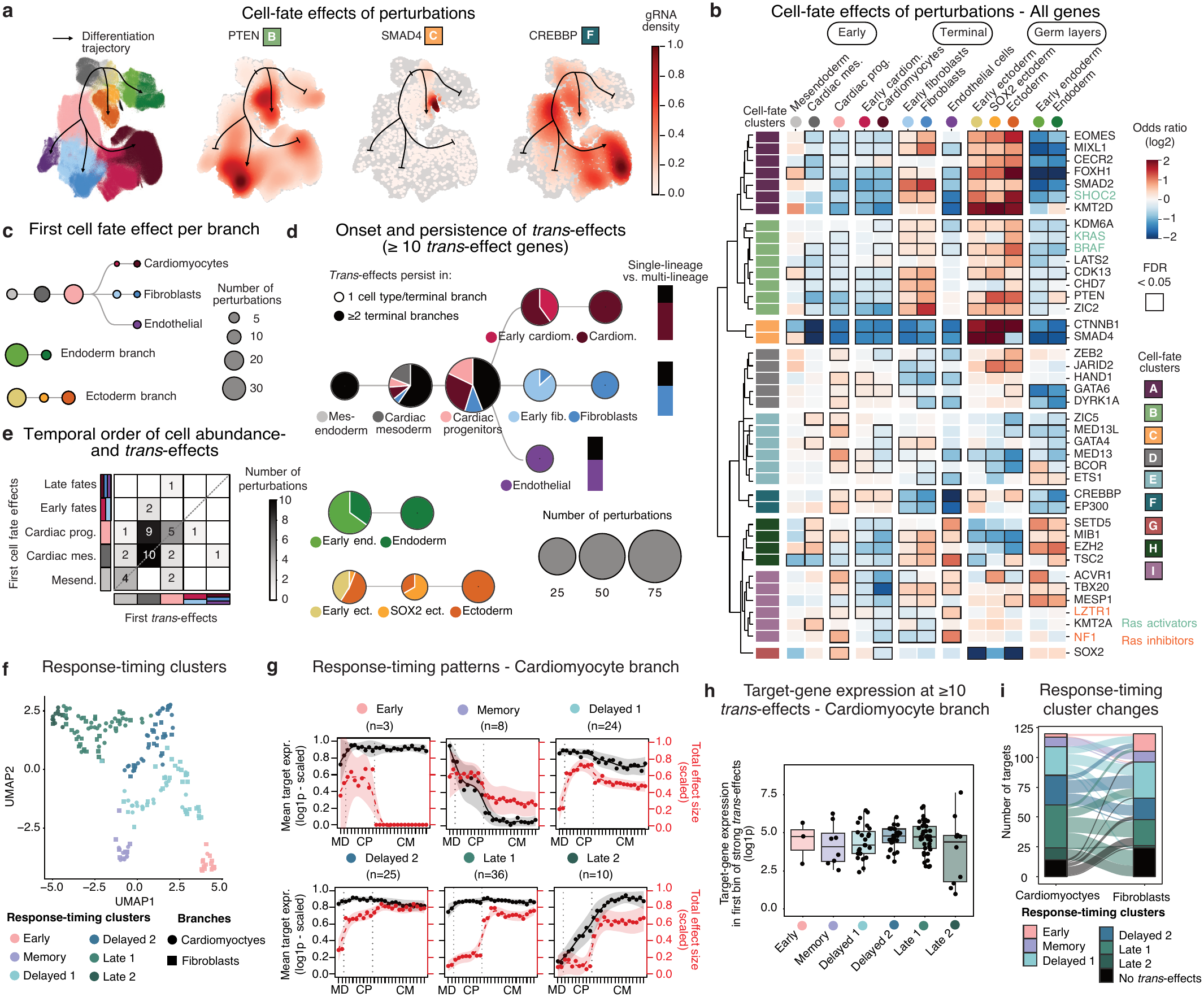
Perturbation-induced cell-fate changes and distinct temporal dynamics between target-gene expression and perturbation responses. a, UMAP embedding of cells from all three integrated datasets, coloured by cell-type annotation (left) or by cell density for selected perturbations (right). Cell-fate clusters are indicated above. **b,** Cell-fate abundance changes for 41 perturbations with FDR < 10LL and absolute odds ratio (log2) ≥ 0.5 in at least two cell types. Colour, odds ratio (log2, capped at ±2) versus negative control (NC) cells (two-sided Cochran-Mantel-Haenszel test). Rows, perturbations grouped into cell-fate clusters A–I; columns, cell types. Black boxes, FDR < 0.05. Teal and orange, Ras activators and inhibitors. **c,** Onset of cell-fate abundance effects. For each of the 41 perturbations, the first developmental stage with a significant abundance change (FDR < 10LL) is shown along each branch. Node size, number of perturbations; node colour, cell type. **d,** Onset and persistence of *trans*-effects across cell types. Dot size indicates the first developmental stage with significant *trans*-effects along each branch. Pie charts denote the fraction of perturbations whose *trans*-effects persist (≥ 10 *trans*-effects, FDR < 0.1) into later cell types, coloured by the most differentiated cell type affected. Bar plots (right) show the fraction of perturbations with first *trans*-effects in terminal cardiac fates, distinguishing single-lineage versus multi-lineage effects. **e,** Temporal order of first cell abundance-and *trans*-effects for the cardiac branch shown in a heatmap. **f,** Joint UMAP of perturbations of concatenated max-scaled target-gene expression and max-scaled total effect size along pseudotime bins, coloured by response-timing clusters. Each point represents one perturbation in the cardiomyocyte (circles) or fibroblast (squares) branch. **g**, Response-timing patterns shown for the cardiomyocyte branch. For each response-timing cluster, mean max-scaled target-gene expression (log1p; black, left axis) and mean max-scaled total effect size (root sum of squared log2 fold changes; red, right axis) across member perturbations are shown per pseudotime bin, from mesoderm (MD) through cardiac progenitors (CP) to cardiomyocytes (CM). Points: bin means; lines: spline fits with shaded 95% confidence intervals; n: number of perturbations per cluster. **h**, Target-gene expression at the onset of strong *trans*-effects (≥10) per response-timing cluster in the cardiomyocyte branch. **i,** Number of perturbations per response-timing cluster in the cardiomyocyte (left) and fibroblast (right) branches (bars), and reassignment of perturbations to different response-timing clusters between branches (flows).

Perturbations in clusters A-C shifted early lineage allocation from cardiac fates and endoderm towards amniotic ectoderm. These cell-fate clusters predominantly depleted cardiac mesoderm, cardiac progenitors, cardiomyocytes, endothelial cells and endoderm while enriching non-neuronal amniotic ectoderm. Cell-fate clusters A and B showed enrichment of fibroblasts, while cell-fate cluster C showed the opposite pattern with depletion of fibroblasts. These clusters included regulators of WNT (*CTNNB1*), PI3K (*PTEN*), Nodal/Activin (*FOXH1*), and Ras/MAPK (*BRAF*, *KRAS*, *SHOC2*) signalling and cardiac mesoderm determinants (*EOMES*, *MIXL1*, *SMAD4* and *ZIC2*)^42^, recapitulating early amniotic ectoderm divergence and gastrulation-dependent mesendoderm specification *in vivo*^45^. This highlights cardiac mesoderm as a sensitive developmental state, potentially linking disrupted cardiac development with the extra-cardiac abnormalities characteristic of syndromic CHD^26^. Many genes in these clusters are implicated in CHD, including *de novo* variants in *PTEN* and *CTNNB1*^36^, as well as *SMAD2*, *CHD7*, *BRAF*, *KDM6A*, *CDK13*, *KMT2D*, *KRAS* and *FOXH1*^41^.

Perturbations in cell-fate clusters D-F acted later in lineage allocation, mainly enriching cardiac progenitors while depleting fibroblast, endothelial and endoderm populations. These clusters included established CHD genes (*BCOR*, *CREBBP*, *GATA6*, *HAND1*, *MED13L*, *ZEB2*)^41^ alongside genes with more limited disease evidence, such as *DYRK1A* and *MED13*, nominating them as CHD-relevant regulators of cardiac lineage allocation. Cell-fate clusters G-I primarily depleted cardiomyocytes while enriching endothelial and endoderm populations, suggesting defects spanning terminal differentiation and germ-layer patterning.

Strikingly, the great majority of the 41 perturbations induced abundance effects that were already detectable in early progenitor states (Fig. 2c). Thus, many CHD-associated genes act during germ-layer formation and early cardiac lineage specification, well before mature cardiac structures emerge. Perturbations of these transient progenitors persisted into downstream lineages, supporting early developmental dysregulation as a common mechanism linking cardiac and extracardiac manifestations of syndromic CHD.

### *Trans*-effects co-occur with or precede cell type abundance changes

Cell-fate abundance captures only one axis of a perturbation’s phenotype. Perturbations can also alter transcriptional programmes within a cell type, either before any detectable change in its abundance or in the absence of one. We therefore asked how transcriptional changes (*trans*-effects) of individual perturbations relate in time to the observed abundance changes. Differential expression testing within each cell type (DESeq2^46^ on pseudobulk replicates per gRNA and differentiation, mean ∼170 cells per perturbation and cell type; Methods) identified significant *trans*-effects (FDR < 0.1) for 386 of 389 tested perturbations in at least one cell type (Extended Data Fig. 3d-g). *P* values were well calibrated, as cells with control gRNAs tested as mock perturbations followed the expected null distribution (Extended Data Fig. 3h). The number of *trans*-effects (mean ∼64 per perturbation and cell type) increased with cell number (Extended Data Fig. 3i), and we focused downstream analyses on 213 perturbations with ≥10 *trans*-effects in at least one cell type.

The majority of these perturbations (134/213) affected cardiac-lineage cells already in mesendoderm, cardiac mesoderm and cardiac progenitors, before terminal fate decisions (Fig. 2d, left). About two-thirds of perturbations with effects in cardiac mesoderm, and almost a third of those affecting cardiac progenitors, subsequently altered more than one terminal fate (cardiomyocytes, fibroblasts and endothelial cells), while effects confined to a single terminal state were less common at these earlier stages. A separate set of perturbations produced no detectable effect until the terminal states were reached, and more than half of these late-onset effects were specific to the cell type in which they first appeared (Fig. 2d, right).

To resolve the temporal order of cell abundance-and *trans*-effects following each perturbation, we compared the earliest developmental stage along the cardiac branch at which *trans*-effects and cell-fate abundance changes were detected (Fig. 2e). This analysis revealed that for most perturbations (35 out of 41), *trans*-effects arose at the same stage as, or before, the abundance change.

### Perturbation effects track cell-state dependency independent of target-gene expression levels

While most *trans*-effects emerged early in differentiation, some appeared only later, raising the question of how their timing and magnitude relate to the target gene’s own expression dynamics. For each perturbation, we estimated time-resolved dynamics of target-gene expression (max-scaled) and total effect size (max-scaled root sum of squared log-fold changes) along the pseudotemporal ordering of the cardiomyocyte and fibroblast trajectories (n=106 perturbed genes with sufficient coverage; Methods; Supplementary Table 9). Hierarchical clustering of the resulting profiles identified six response-timing clusters (Fig. 2f, g; Extended Data Fig. 4a), which can be broadly grouped into (i) an early-effect response-timing cluster, in which *trans*-effects were restricted to mesoderm and cardiac progenitors; (ii) a memory response-timing cluster, in which *trans*-effects persisted as target-gene expression declined; and (iii) delayed/late-effect response-timing clusters, in which *trans*-effects emerged only at later stages, despite target-gene expression.

Target genes were ubiquitously expressed in all response-timing clusters except Late 2 and Memory (low expression at early and late pseudotime, respectively), whereas their *trans*-effects followed distinct temporal patterns. In the Memory cluster, *trans*-effects persisted after the target gene was no longer expressed. This cluster contained a higher proportion of transcriptional regulators than the other clusters (Extended Data Fig. 4b), consistent with the expectation for TF perturbations to have long-lasting downstream consequences. For all clusters, gene expression levels of the perturbed gene were high at the onset of *trans*-effects (Fig. 2h, Extended Data Fig. 4c), indicative of a critical regulatory role. Finally, when the same perturbations were compared between the cardiomyocyte and fibroblast branches, perturbations classified as Early and Memory largely retained their response timing, whereas perturbations classified as Delayed frequently changed cluster and Late perturbations often lacked *trans*-effects in the other branch (Fig. 2i, Extended Data Fig. 4d), indicating that the timing of later effects depends on lineage context.

Collectively, these data suggest that target-gene expression is necessary but not sufficient to detect *trans*-effects. Beyond requiring expression, the onset and persistence of *trans*-effects are largely decoupled from the target gene’s own expression dynamics, reflecting instead the point at which a given cell state becomes functionally dependent on the target gene.

### Consensus GRNs uncover hierarchical TF control and regulators of cardiac lineage specification

scPerturb-seq provides a powerful approach for reconstructing GRNs by systematically perturbing TFs and measuring their *trans*-effects. The snMultiome-seq component of our atlas offers a complementary route to GRNs: it captures accessible chromatin regions, enabling inference of TF binding, but does not measure the downstream consequences of TF perturbation. To compare and combine both approaches to derive consensus GRNs, we inferred cell-type-specific TF-enhancer-gene GRNs from matched, temporally resolved snMultiome-seq data. To construct cell-type-specific enhancer-mediated gene regulatory networks (eGRNs), we first fitted scDORI^47^ and then integrated its (i) predictions of TF binding at enhancers^15^, and (ii) enhancer-to-gene links with (iii) cell-type-specific enhancer accessibility (Fig. 3a; Methods).

**Figure 3.**
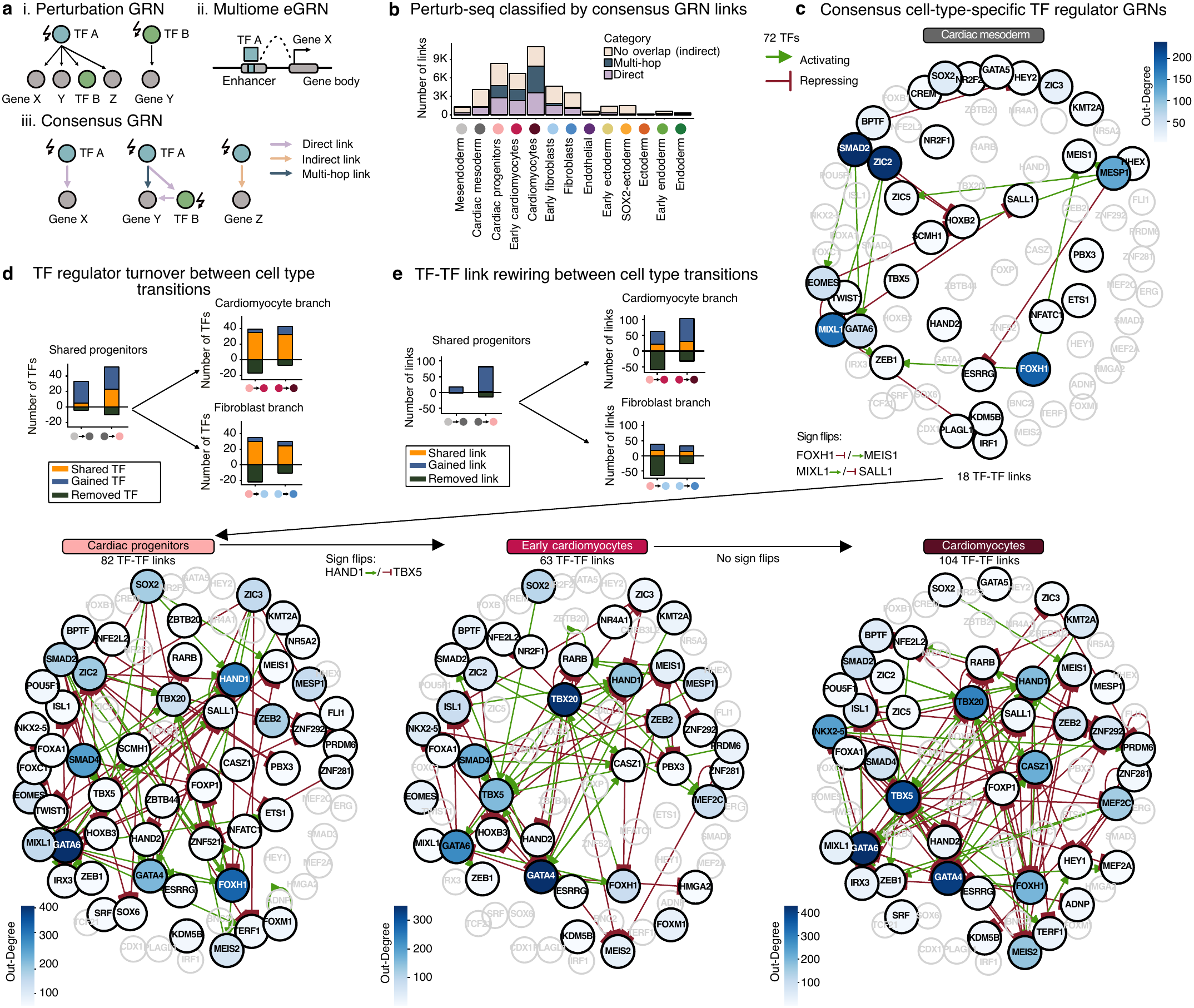
Consensus gene regulatory networks from combined scPerturb-seq and snMultiome data. a, Approach for classifying the mode of TF regulation by combining perturbation-and multiome-derived GRNs. Shown are the GRN features captured by scPerturb-seq *trans*-effects (i) versus enhancer-mediated GRNs (eGRN) estimates from snMultiome-seq (ii), and the resulting classification of regulatory links in consensus GRNs as direct, multi-hop or indirect (iii). **b,** Stacked barplot of perturbation-derived regulatory links, classified according to the approach in **(a)**. **c,** TF-TF consensus GRNs per cell type along the cardiomyocyte branch, ordered by differentiation. Nodes, TFs regulators coloured by out-degree (number of downstream genes); edges, activating (green) or repressing (red) TF-TF links; faded grey, TFs connected only in other cell types; labels, links that switch sign between consecutive cell types (TF-only consensus GRN; 72 TFs with consensus links in the cardiomyocyte or fibroblast branch). **d, e,** Rewiring of TF-TF networks between consecutive cell types, from shared progenitors into the cardiomyocyte (top) and fibroblast (bottom) branches. Bars show TF regulators **(d)** or TF-TF links **(e)** that are shared, gained or lost at each transition, ordered by differentiation (left to right). Shared progenitors: mesendoderm, cardiac mesoderm and cardiac progenitors; includes TFs with out-degree > 1; x-axis dots, cell types of each transition, coloured as in **(b)**.

Intersecting multiome-derived eGRNs with perturbation-derived links revealed significant overlap in key cardiac cell types (Fig. 3b, Extended Data Fig. 5a-c; Supplementary Table 10), however, many perturbation-derived links had no direct counterpart in the multiome-derived eGRN. This is partly expected, because perturbations capture both direct and downstream effects, whereas eGRN links represent direct TF binding. Indeed, TF perturbations frequently altered expression of genes lacking a direct TF-enhancer-gene link, consistent with multi-step cascades (TF-A → TF-B → Gene K), allowing us to map 9% (SOX2-ectoderm) to 71% (cardiomyocytes) of perturbation-derived TF-gene links onto multiome-derived GRNs (Fig. 3b, Extended Data Fig. 5c,d). We used regulatory relationships supported by both assays to define a consensus GRN with a particularly stringent set of links for downstream interpretation.

As expected for cell-type-specifying master regulators, the TFs with the most downstream targets (out-degree) in each cell type included central regulators of cardiac mesoderm (*ZIC2*, *SMAD2*, *MESP1*, *MIXL1*, *FOXH1*, *EOMES*), early cardiomyocytes (*TBX20*) and cardiomyocytes (*CASZ1*, *TBX5*, *NKX2-5*) (Fig. 3c, Extended Data Fig. 5e, f). In fibroblasts, *ETS1* was among the most connected TFs, consistent with its role in epithelial-to-mesenchymal transition and its association with hypoplastic left heart syndrome^48,49^. GRN TF regulators were largely conserved across closely related cell types, such as early and mature cardiomyocytes, where minimal rewiring is expected, validating our approach for defining consensus GRNs (Extended Data Fig. 5g).

We next investigated the extent to which TF regulators change across cell differentiation. Within the mesoderm branch, we observed larger TF regulator turnover (fewer shared TFs) early in differentiation, whereas they stabilised from the cardiac progenitor state onward (Fig. 3d). In contrast, the links from these regulators to their downstream TFs (TF-TF links) were rewired more extensively and continued to change throughout differentiation (Fig. 3e). These patterns of stable TF regulators in later differentiation stages were also observed in solely perturbation-and multiome-derived networks (Extended Data Fig. 6a-d).

TFs in the consensus GRNs showed both repressive and activating effects, depending on the downstream gene, cellular context and developmental stage^50^. We therefore asked whether individual TFs showed a consistent bias towards repressive or activating activity across cell types (Extended Data Fig. 6e). Most perturbed TFs showed a predominantly repressive signature, with few exceptions (*NFE2L2*, *ZNF292*, *RARB*). This suggests that mesoderm differentiation relies largely on TF-mediated repression of alternative programmes, which, together with a few activating lineage regulators, stabilises terminal cell identity^51,52^.

The consensus GRNs also placed *HOXB2* and *HOXB3* under cardiac transcriptional control. In cardiac mesoderm, *ZIC2*, *SMAD2* and *EOMES* repressed *HOXB2* (Fig. 3c), and its suppression might drive the pronounced abundance changes observed upon perturbation of these three factors (Fig. 2b). Although *HOXB2* has been detected in a distinct Mesp1L second heart field cardiovascular progenitor subpopulation in mice^53,54^, neither *HOXB2* nor *HOXB3* has been assigned upstream regulators in a cardiac GRN, or tested functionally in cardiac lineage specification. *HOXB3* was likewise repressed, by *SMAD2*, *MIXL1* and *ZIC2* in cardiac progenitors and by *GATA4*, *TBX20*, *ZIC2* and *SMAD4* in early cardiomyocytes. These findings expand the set of cardiac HOX regulators beyond *HOXA1*, *HOXB1*, *HOXB5* and *HOXB6* to include *HOXB2* and *HOXB3* as candidate regulators of lineage specification^53^. Beyond nominating individual regulators, this multiome-and perturbation-anchored consensus network offers a mechanistic lens onto a recurring pattern in our data: perturbations of different genes often produce strikingly similar transcriptional signatures.

### Shared transcriptional responses reveal co-regulator networks across cell types

Genes acting in the same regulatory network, pathway or protein complex often induce similar *trans*-effects when perturbed^55^. We therefore clustered perturbation–cell type combinations by *trans*-effect similarity (Pearson correlation of z-scored log-fold changes; perturbation–cell type combinations with ≥10 significant *trans*-effects), identifying 50 co-regulator clusters (Fig. 4a; Extended Data Fig. 7a; Supplementary Tables 11 and 12; Methods). After discarding clusters with weakly correlated, unspecific *trans*-effects (Methods), 48 co-regulator clusters comprising 125 unique perturbations remained (Extended Data Fig. 7b). Among these, we identified both cell-type-specific and-mixed co-regulator clusters, indicating that co-regulatory programmes^56^ can either be restricted or shared across cell identities, respectively (Fig. 4b).

**Figure 4.**
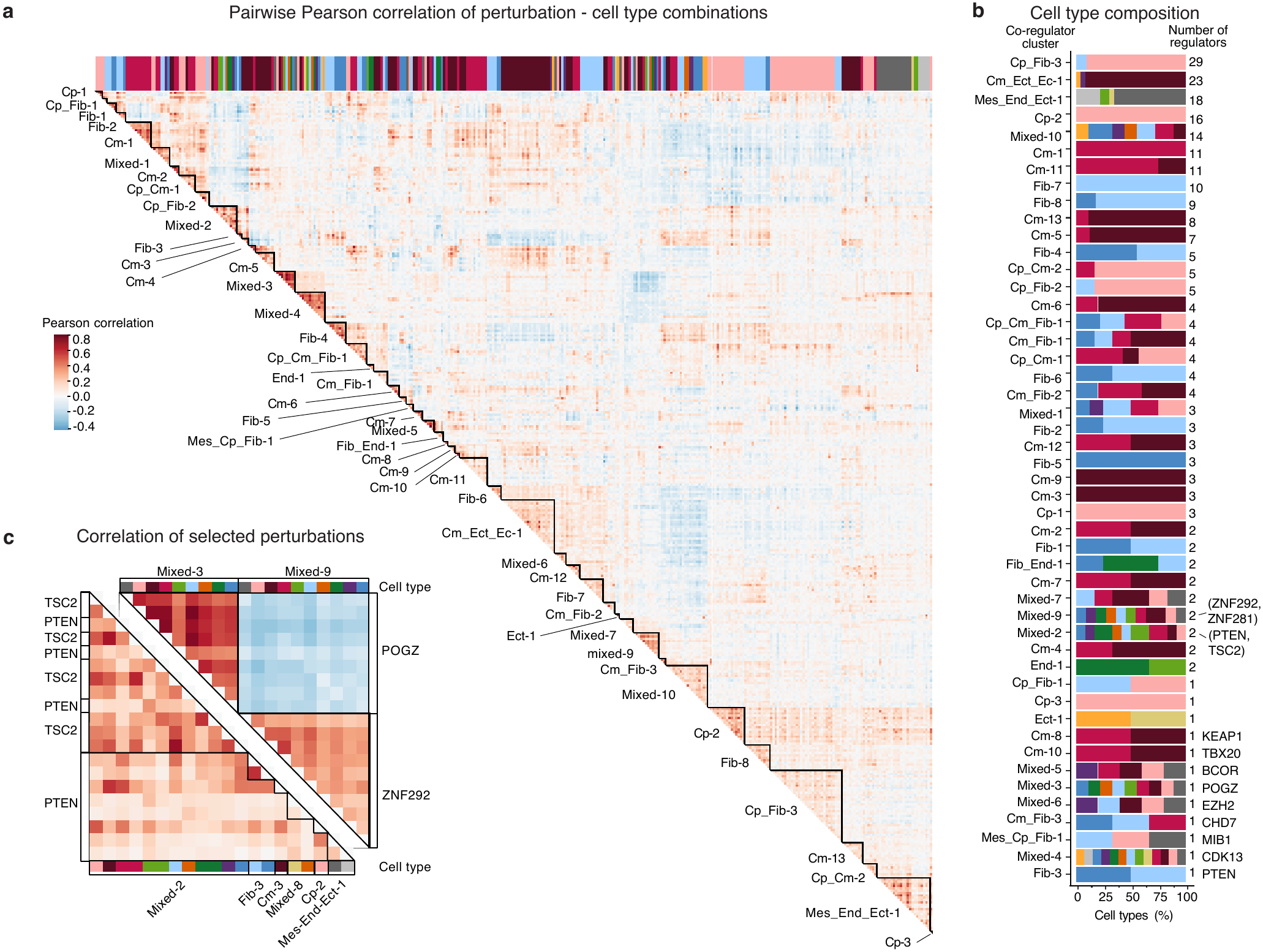
Co-regulation within and across cell types. a, Pairwise similarity of transcriptional responses between regulator-cell type pairs, grouped into co-regulator clusters by hierarchical clustering (k = 50 clusters; regulator-cell type pairs with ≥ 10 *trans*-effects). Colour, Pearson correlation of *trans*-effect profiles (z-scored log-fold changes). Two clusters with weakly correlated, unspecific *trans*-effects are shown in Extended Data Fig. 7b. **b,** Cell type proportions for each of the 48 co-regulator clusters. **c,** Zoom-in view highlighting selected regulators, revealing a strong, previously unreported anticorrelation between *POGZ* and *ZNF292* (upper triangle) and a positive correlation between *PTEN* and *TSC2* (lower triangle), consistent with their established roles in a shared pathway. Column labels denote co-regulator clusters; colour bars indicate cell types.

Several identified co-regulator clusters captured perturbations in the same protein complex (e.g., *MED13/MED13L)* or pathway (e.g., *PTEN/TSC2)* indicating that this approach can capture known regulatory relationships (Fig. 4c, Extended Data Fig. 7c), similar to analysis conducted in perturbation-screens of individual cell types^56^. We also identified clusters of perturbations with anticorrelated *trans*-effects, most strongly between zinc-finger regulators *ZNF292 (*Mixed-9) and *POGZ* (Mixed-3), for which no direct interaction has been reported (Fig. 4c). Both are high-confidence autism genes, suggesting convergence on a common neurodevelopmental programme, while our data suggest an additional regulatory role in heart development^57,58^. These results reveal shared and antagonistic *trans*-modules co-regulated by distinct sets of perturbations.

### Cell-fate and co-regulatory perturbations converge on CHD

Having examined perturbations through various lenses, we next asked if perturbations that exhibit strong *trans*-effects and co-regulatory effects converge on disease-relevant genes. We found that co-regulator perturbations, or more generally, perturbations with ≥10 *trans*-effects in at least one cell type were significantly enriched for CHD genes (Extended Data Fig. 7d). Perturbations within individual cell-fate clusters overlapped with distinct co-regulator clusters, indicating that perturbations driving abundance changes also converge on shared transcriptional programmes (Extended Data Fig. 7e). Similar overlap was also observed when comparing co-regulators and cell-fate effect clusters to response-timing clusters (Extended Data Fig. 7f, g). Thus, disease-relevant perturbations reshape cardioid differentiation along two coupled dimensions: they alter which cell fates are reached and reprogram the molecular identity of the cells that emerge.

### Differentiation reshapes early perturbation signatures into disease-relevant programmes

Co-regulators have previously been identified within individual cell types^55^, but whether co-regulatory relationships persist through differentiation has not been systematically assessed. Many CHD genes already act in cardiac progenitors, yet structural defects emerge only as cells differentiate. We therefore asked whether co-regulators in cardiac progenitors remain co-regulators and whether they regulate the same or different sets of genes in differentiated cell types. Notably, only 34 of the 64 co-regulators identified in cardiac progenitors retained co-regulation throughout differentiation with at least one of their original progenitor co-regulators (Fig. 5a, b).

**Figure 5.**
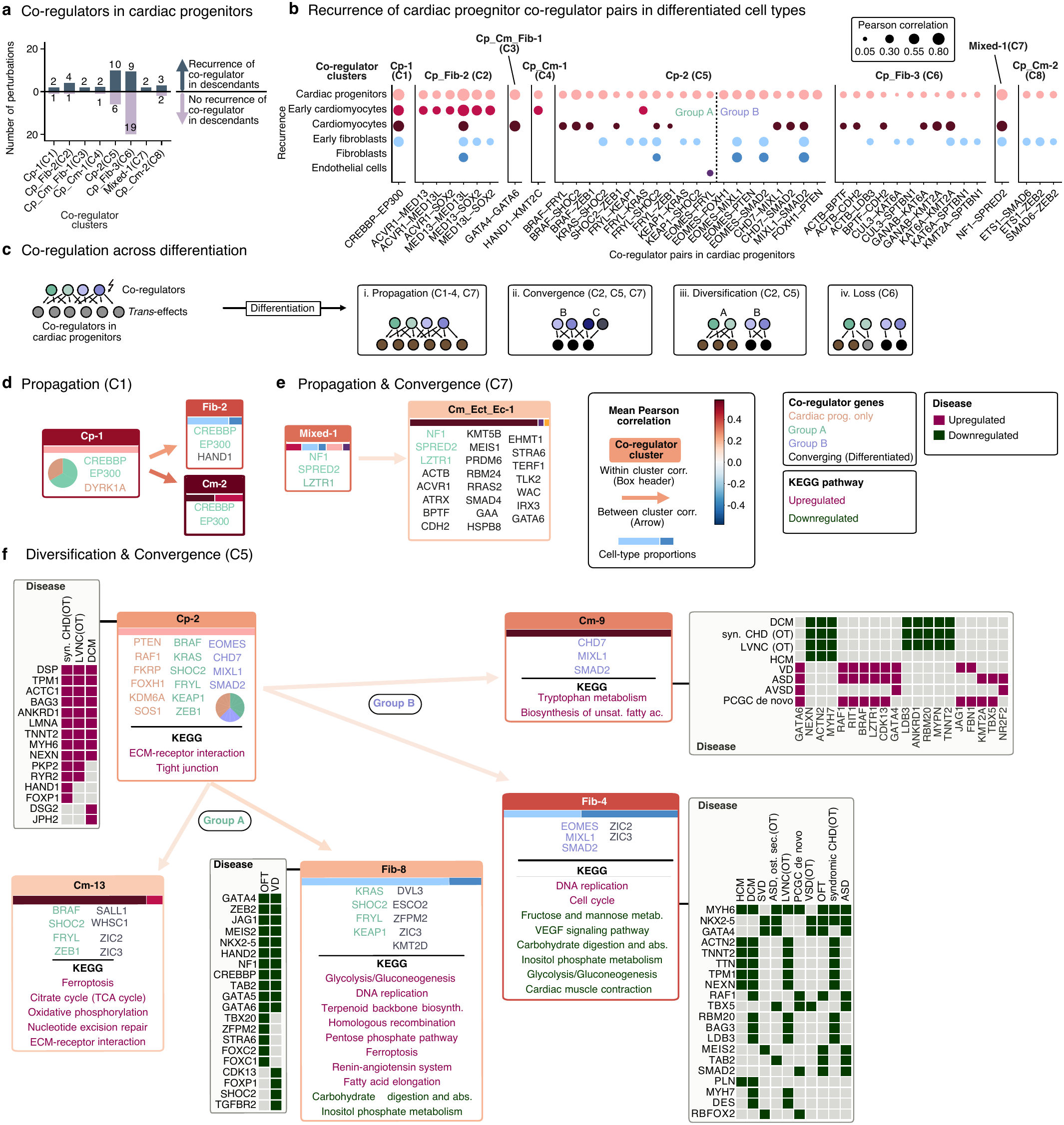
Propagation, diversification, and convergence of co-regulators during cardiac differentiation. a, Number of cardiac progenitor co-regulators that recur (dark blue, above axis) or do not recur (light purple, below axis) with at least one co-regulator in a differentiated cell type, grouped by the eight cardiac progenitor-containing co-regulator clusters. Corresponding cluster families (C1-C8) from panel **(c)** are indicated. **b,** Pearson correlations of perturbation pairs identified in cardiac progenitor co-regulator clusters and retained in at least one co-regulator cluster of differentiated cell types. Pairs are grouped by their cardiac progenitor co-regulator cluster. Dot size denotes Pearson correlation of *trans*-effects across the indicated cell types. **c,** Perturbations with correlated transcriptional responses (co-regulators) in cardiac progenitors can remain correlated upon differentiation (propagation), split into distinct groups (diversification), converge from previously distinct groups (convergence), or lose coordination (loss). **d-f,** Representative co-regulator cluster families showing *trans*-effect correlations (header colour: mean within-cluster correlation; arrow colour: mean between-cluster correlation), KEGG pathway and disease enrichment for CHD and cardiomyopathy (magenta: upregulated; green: downregulated), and cell-type proportions within clusters (stacked bars). Propagation in co-regulator family C1 **(d)**: *CREBBP*, *EP300* and *DYRK1A* perturbations show a coordinated response in cardiac progenitors (Cp-1) that persists into cardiomyocytes (Cm-2) and fibroblasts (Fib-2). Propagation and convergence in co-regulator family C7 **(e)**: Mixed-1 co-regulator cluster (*NF1*, *SPRED2*, *LZTR1*) stays coordinated from a mixed progenitor state into Cm-Ect-Ec-1 and converges with additional regulators. Diversification and convergence in co-regulator family C5 **(f)**: Co-regulators in cardiac progenitors (Cp-2) split upon differentiation into group A (*BRAF*, *KRAS*, *SHOC2* and interacting TFs *FRYL*, *KEAP1*, *ZEB1*), giving rise to Cm-13 and Fib-8; and group B (*EOMES*, *CHD7*, *MIXL1*, *SMAD2*), remaining coordinated in Cm-9 and Fib-4; *ZIC2*/*ZIC3* converge with group A in Cm-13 but with group B in Fib-4.

To identify more general patterns of this rewiring, we grouped co-regulator clusters into families based on the regulators they have in common (seven co-regulator families after discarding weakly overlapping cluster C8; Extended Data Fig. 8a-c; Methods). Inspection of these co-regulator families (C1-C7) revealed four modes in which a co-regulator cluster identified in cardiac progenitors can evolve during differentiation (Fig. 5c, Extended Data Fig. 8a-c): (i) Propagation, in which the progenitor co-regulator cluster remains intact; (ii) Diversification, in which the cluster splits into sub-clusters that remain co-regulated internally but no longer co-regulate with each other; (iii) Convergence, in which the cluster is retained and recruits additional regulators; and (iv) Loss, in which cluster members no longer co-regulate. These modes were not mutually exclusive, and some co-regulator families combined several of them.

Propagation was the most common mode (five of seven co-regulator families: C1-C4 and C7), but conserved co-regulation did not imply conserved function: co-regulators maintained correlated transcriptional responses across differentiation even though the *trans*-effects themselves changed (Fig. 5d, e, Extended Data Fig. 9a-d, Extended Data Fig. 10a). For example, CREBBP and EP300 maintained co-regulated *trans*-effects from cardiac progenitors (Cp-1) to cardiomyocytes (Cm-2) and fibroblasts (Fib-2) (Fig. 5d, Extended Data Fig. 9a), suggesting inheritance of a shared progenitor state or sustained regulatory coupling. In cardiac progenitors, their loss suppressed focal adhesion (Extended Data Fig. 9a, Supplementary Tables 13–16) and was associated with prolonged progenitor-like states (Fig. 2b, cell-fate cluster F). In cardiomyocytes, loss of CREBBP and EP300 instead suppressed excitation-contraction coupling (calcium) and metabolic maturation pathways (PI3K-AKT, HIF-1 signalling, oxidative phosphorylation), downregulating cardiomyopathy (DCM, HCM, left ventricular non-compaction – LVNC) and syndromic CHD genes, while upregulating CHD genes (ASD, valve defects – VD, PCGC *de novo*) involved in RAS/MAPK signalling and chromatin regulation (Extended Data Fig. 9a). Thus, co-regulatory relationships between perturbations can persist across differentiation while their downstream programmes shift, including opposing regulation of CHD-and cardiomyopathy-associated genes, potentially influenced by residual cardiomyocyte immaturity.

Conversely, similar progenitor responses did not predict similar responses after differentiation. In co-regulator family C5, perturbations highly correlated in progenitors (Cp-2) split into two distinct groups after differentiation (Fig. 5f; Extended Data Fig. 8c). Group A, comprising Ras/MAPK regulators (*BRAF*, *KRAS*, *SHOC2*) and interacting transcriptional regulators (*FRYL*, *KEAP1*, *ZEB1*), gave rise to Cm-13 and Fib-8. In contrast, group B, comprising early transcriptional regulators (*EOMES*, *CHD7*, *MIXL1*, *SMAD2*), remained co-regulators in Cm-9 and Fib-4. Despite this shared progenitor signature, the resulting cardiomyocyte clusters (Cm-9 and Cm-13) shared no co-regulators, and their *trans*-effects were mildly anticorrelated (Extended Data Fig. 8c). In addition to diversification, *ZIC2* and *ZIC3* converged with group A in cardiomyocytes (Cm-13) but with group B in fibroblasts (Fib-4). Thus, similar progenitor responses can resolve into distinct lineage-dependent transcriptional programmes, detectable through time-resolved data.

Loss of coordination, exemplified by Cp-Fib-3 (C6), did not imply loss of function: co-regulators weakly correlated in progenitors showed no coordination in terminal states despite substantial individual *trans*-effects (Extended Data Fig. 10b). This suggests that differentiation can erase coordination established in cardiac progenitors. In contrast, other perturbations exhibited markedly fewer detectable *trans*-effects after differentiation, consistent with attenuation of their early transcriptional signatures (Extended Data Fig. 10c, d).

Ras/MAPK pathway perturbations further revealed contrasting regulatory responses associated with cardiac disease genes. Perturbations of Ras activators (*BRAF*, *KRAS*, *SHOC2*; C5) and Ras inhibitors (*NF1*, *SPRED2*, *LZTR1*; C7) showed anticorrelated *trans*-effects in progenitor-containing co-regulator clusters (Cp-2 and Mixed-1; Fig. 5e, f, Extended Data Fig. 8b), accompanied by opposing CHD and cardiomyopathy gene set enrichment. Furthermore, they had opposing effects on cell-type abundance (Fig. 2b), although their metabolic pathway signatures were not consistently opposing (Fig. 5e, f; Extended Data Fig. 9d). Perturbations in co-regulator family C2 correlated positively with Ras inhibitors (C7) and negatively with Ras activators (C5) (Extended Data Fig. 8a-c). Loss of Ras inhibitors mimics gain-of-function mutations in Ras activators, which have been implicated in CHD^34,59–61^. Together, these data implicate Ras signalling in both cell-fate and disease-associated transcriptional programmes, linking altered cardiac differentiation to CHD and cardiomyopathy.

Together, these findings show that differentiation reshapes the transcriptional consequences of gene perturbations, including CHD-associated genes, through propagation, diversification, convergence or loss, acting as an active transformative layer between genotype and phenotype. Conserved co-regulatory relationships can acquire distinct downstream functions, similar progenitor responses can diverge upon differentiation, and loss of co-regulation does not imply loss of individual regulator activity. Opposing Ras pathway perturbations link signalling activity to cell fate and disease-associated programmes. Our data further reveal a developmental shift in disease-associated transcriptional responses, from morphogenetic programmes linked to structural CHD in progenitors towards contractile and metabolic programmes associated with cardiomyopathy in cardiomyocytes, suggesting that CHD genes can have temporally distinct pathogenic consequences.

## Discussion

In this study, we established a time-resolved cardioid scPerturb-seq screen and combined it with multiomic profiling in cardioids. We mapped how perturbations affect cell fate, transcriptional state and regulatory circuitry throughout cardiac differentiation. Cardioids provide experimental access to a window of human cardiogenesis that current *in vivo* data lack, as these datasets profile from around 28–31 dpf onwards^4–14^. Cardiac mesoderm and the first cardiac progenitors are specified in the anterior lateral mesoderm around 18–20 dpf, while bilateral heart fields fuse into a contracting linear heart tube by approximately 22 dpf^23^. Profiling of CHD specimens is performed later, typically following infant or paediatric surgery^13^. The dense temporal sampling design employed in this study takes a crucial step towards closing this gap, identifying early cardiac development as a sensitive period for CHD-associated genes. During this window, perturbations of patterning pathways, chromatin regulators and TFs shifted the balance between germ layers towards an ectoderm-like fate, potentially contributing to extracardiac manifestations of syndromic CHD^26^. Genes with similar abundance effects frequently converged on shared co-regulated *trans-*effects within cell states, tying fate and molecular phenotypes together.

Cardiac mesoderm emerged as a particularly critical cell state for CHD-associated genes, with key disease genes such as *ZIC2*, *SMAD2*, *EOMES*, *MIXL1* and *FOXH1* among the strongest drivers of fate reallocation. This is consistent with mouse studies, showing that *Mesp1*^+^ mesoderm segregates into cardiovascular lineages within hours of gastrulation^54^, and that chromatin regulators acting at primed cardiac enhancers canalise mesoderm differentiation^62^. Our data also provide new insights into the factors that govern the timing of *trans*-effects, with functional dependency rather than expression level of the target being the key determinant.

Sampling differentiation densely rather than at a single endpoint made the temporal order of perturbation effects accessible. *Trans*-effects arose at the same stage as, or before, the corresponding abundance change. Furthermore, the timing of *trans*-effects was largely decoupled from the target gene’s own expression dynamics, persisting after expression declined, particularly for TFs, or emerging only after a delay. Expression is therefore necessary but not sufficient; when a perturbation has downstream effects reflects the point at which a cell state becomes functionally dependent on the gene. This limits what observational *in vivo* data can establish about which cell types are functionally affected by disease-associated genes^4–14^, as trait-associated genes may be overlooked in mature cell types where they are no longer expressed, despite persistent effects of their activity in progenitors. Pairing the screen with enhancer-driven eGRNs from snMultiome-seq addressed a different limitation: perturbation effects alone cannot separate direct regulation from multi-step cascades (TF-A → TF-B → Gene K), whereas enhancer binding alone does not reveal the consequences of removing a TF. Combining both yielded consensus GRNs which nominated *HOXB2* and *HOXB3* as candidate cardiac regulators.

Early *trans*-effects did not simply persist through differentiation. Co-regulatory relationships in progenitors were propagated, diversified, converged or lost as cells committed to cardiomyocyte or fibroblast fates, and even persisting relationships controlled different programmes in their descendants. Together with the continued rewiring of TF-TF links after the set of TF regulators had stabilised, this suggests that regulatory modules are reassembled from a largely stable set of regulators as cells differentiate. Co-regulator maps from single cell states^56^ may therefore capture only snapshots of context-dependent relationships, a dependence that loss of *Hand2* in mice illustrates in vivo^63^.

Although cardioids provide access to early human cardiogenesis, they lack the full anatomical complexity, the mechanical and hemodynamic inputs, and the later-emerging cell types of the *in vivo* heart. The protocol favours ventricular lineages and underrepresents some heart-field derivatives, but remains well suited to capture early gastrulation-like fate decisions across mesoderm-, endoderm-and ectoderm-like states. Links to CHD therefore rest on gene-set enrichment and fate shifts rather than structural phenotypes. Furthermore, perturbed cells developed in mosaic cardioids, where neighbours may mask non-cell-autonomous effects. Power to detect *trans*-effects depended on cell number, so absence of effects in sparser populations should be interpreted cautiously. In this study, we assayed loss of function only; dosage effects of key TFs have been implicated in CHD and GRN rewiring and would require an allelic-series design^64^.

This study provides a framework to dissect developmental gene function with temporal and cellular resolution, linking disease genes to specific developmental windows and transcriptional programmes. We showed that early human cardiogenesis is shaped by dynamic regulatory dependencies that static or current *in vivo* data cannot resolve. By jointly profiling gene expression, chromatin accessibility and perturbation responses over time, our data capture context-dependent, temporally changing perturbation effects that are largely absent from existing resources. These data provide a foundation for developing and benchmarking predictive models that infer GRNs, predict perturbation effects and anticipate cell-fate decisions, and for advancing efforts towards virtual cell models.

## Supplementary Tables

|  |  |
|---|---|
| Table 1 | Perturbed genes and gRNA information |
| Table 2 | Thresholds for downstream data pre-processing |
| Table 3 | Number of cells per dataset and cell type |
| Table 4 | Marker genes used for cell type annotation |
| Table 5 | Disease gene list and gene expression enrichment |
| Table 6 | Perturbed gene selection for perturbing genes |
| Table 7 | Non-targeting gRNA information |
| Table 8 | Cell type abundance changes and cell-fate clusters |
| Table 9 | Response-timing clusters; Gene expression and total effect size per perturbations and pseudotime-cell type bin |
| Table 10 | Cell-type-specific consensus GRNs |
| Table 11 | Pairwise correlation of *trans*-effects |
| Table 12 | Co-regulator clusters from hierarchical clustering |
| Table 13 | KEGG pathway enrichment in co-regulator clusters |
| Table 14 | GO term enrichment in co-regulator clusters |
| Table 15 | Disease gene list combined from different sources |
| Table 16 | Disease gene enrichment in co-regulator clusters |
| Table 17 | Differentiation experiments and sequencing runs for scPerturb-seq |
| Table 18 | Primer sequences for cloning and genotyping |

## Methods

### Cell culture

WTC-11 iPSCs (Coriell Institute for Medical Research, GM25256) were verified to display a normal karyotype, were contamination free and were regularly tested for mycoplasma. Cells were cultured in Essential 8 medium (E8; Thermo Fisher Scientific, A1517001) on Vitronectin XF-coated (Gibco^Tm^, A400458) tissue culture plates. Cells were maintained at 37L°C and 5% CO_2_. iPS cells were split using Accutase (PAN-Biotech, P10-21500) and E8 supplemented with 10LµM Y-27632 dihydrochloride (RI; Tocris, 1254). After single-cell dissociation, 1 volume of E8L+ 10 µMLRI was added, and cells were spun at 200g (5Lmin), resuspended and plated in E8L+LRI. All experiments involving human cells were conducted in accordance with institutional guidelines and regulations. The parental tissue donor provided written informed consent under protocols approved by the repository’s governing Institutional Review Board, enabling broad genetic data access and public distribution for biomedical research.

### Cardioid generation

Cardioids were grown using an adapted protocol from Hofbauer *et al*.^25^ WTC-11 iPSCs were grown until confluency and treated with Accutase to generate a single-cell suspension. 1 volume of E8 supplemented with 5LµM RI was added to the well, cells centrifuged at 200g and resuspended in 5LµM RI. Cells were seeded at a concentration of 3,000 cells per well in an Ultra-low attachment plate (Corning). Plates were spun at 200g to aid embryonic body formation. Base cardiac differentiation media (CDM) consists of 50% F12 (GIBCO - #31765068), 45% IMDM (GIBCO - #21980065) and 5% Concentrated lipid (1x - GIBCO - #11905031) supplemented with 0.05% (w/v) BSA (Europa Bioproducts - EQBAH-0500), 15 µg/ml Transferrin (Sigma-Aldrich - T3309-100MG), 0.0004% Monothioglycerol (Sigma-Aldrich - M6145) and Penicillin-Streptomycin (Gibco-15276355). Differentiation started 24 h after seeding (considered day 0) by replacing media with 200 µl CDM supplemented with 3 ng/ml FGF2 (R&D Systems - 233-FB-025/CF), 5 µM LY294002 (Tocris - #1130), 4 ng/ml Activin A (R&D Systems - 338-AC-010/CF), 10 ng/ml BMP4 (R&D Systems - RD-314-BP-050) and 4 µM CHIR99021 (R&D Systems - RD-4423/50). After 36–40 h (day 2) media was exchanged to 200 µl CDM supplemented with 8 ng/ml FGF2, 10 ng/ml BMP4, 10 µg/ml insulin (Roche - #11376497001), 2 µM C59 (Tocris - #5148) and 0.05 µM Retinoic acid (Sigma Aldrich - #R2625) and kept until including day 5 (four media changes). On both day 6 and day 7 media was exchanged to 200 µl CDM supplemented with 8 ng/ml FGF2, 10 ng/ml BMP4 and 10 µg/ml insulin. From day 8 onwards media was exchanged every two days to 200 µl CDM supplemented with 10 µg/ml insulin.

### Cardioid sample preparation scRNA-seq, snMultiome-seq and scPerturb-seq

For scRNA-seq and scPerturb-seq datasets, cardioids were transferred to a 1.5 ml microcentrifuge tube, washed with 1 ml PBS (-/-) and 750 µl of dissociation reagent from the STEMdiff cardiomyocyte dissociation kit (Stem Cell Technologies - #05025). Cardioids were incubated at 37 °C for 15 min at a Thermoblock shaking at 1,000 rpm and inverted every 5 min to aid dissociation. Dissociated cell suspension was transferred to a 15 ml tube containing pre-cooled 5 ml of CDM. Cells were spun at 500 g for 5 min in a swinging bucket rotor, supernatant removed and resuspended in 500 µl 1xPBS (-/-). This was followed by cell fixation according to the user guide of the ScaleBio^TM^ Sample Fixation Kit (RevB_18Apr23) and cells were stored at-70 °C. For snMultiome-seq, cardioids were transferred to a 1.5 ml Eppi, washed with 1 ml PBS (-/-), supernatant removed and snap frozen using liquid nitrogen. Snap-frozen cardioids were processed using the Chromium Nuclei Isolation Kit (10x Genomics - PN-1000494) according to the user guide (10x Genomics - CG000505 - Rev A) to extract nuclei. Cells or nuclei were sampled at the indicated time points of cardioid generation (day 1.5 = day 2, day 2.5 = day 3, day 3.5 = day 4, day 4.5 = day 5, day 5.5 = day 6, day 6.5 = day 7, day 7.5 = day 8, day 14.5 = day 15). Overview of sampling time points, number of differentiation experiments for each dataset and sequencing runs can be found in supplements (Extended Data Fig. 1c, Supplementary Table 17).

### Cloning, molecular biology and generation of transgenic iPSCs

For the constitutive Cas9-expressing cell line used for screening, the corresponding transgene was inserted into the *AAVS1* locus in WTC-11 iPSCs as previously described using TALENS^65–67^. The *AAVS1* targeting vector containing the homology arms, the CAGG promoter, eCas9, dTomato, and a WPRE was a kind gift from J. A. Knoblich (Institute of Molecular Biotechnology of the Austrian Academy of Science, Vienna BioCenter). eCas9 was replaced with Cas9 using a ligation-based approach (AgeI/SphI), using pSpCas9(BB)-2A-GFP (Addgene - 48138) as a donor plasmid. To generate transgenic Cas9-expressing iPSCs, WTC-11 cells were electroporated with the corresponding homology plasmid (3 µg per electroporation) and two TALEN plasmids (0.75 µg per electroporation each) targeting the AAVS1 locus (Addgene, 52341 and 52342). iPSCs were dissociated into a single-cell suspension and counted, and 1L×L10^6^ cells were electroporated using the CB-150 program of the 4D-Nucleofector System and the P3 Primary Cell 4D-Nucleofector X Kit L (Lonza, V4XP-3024), according to the manufacturer’s protocol and plated in E8L+LRI. Cells were sorted 7–10Ldays after electroporation for dTomato using a BD Fortessa instrument running Diva (V9.0.1) software, plated at low density in E8L+LRI, grown to colonies, picked and genotyped (Supplementary Table 18). Positively genotyped clones were checked for homogenous dTomato signal and validated for activity in a corresponding assay, and three clones of each cell line were subjected to a genotyping array screening using an Infinium Global Screening Array-24 kit (Illumina, 20030770) to check for chromosomal rearrangements in iPS cell clones. Only clones that showed no or minor differences to the WTC-11 wild-type parental cell line were kept. The CROP-seq construct CROP-EIF2A-Puro-T2A-GFP was generated by Gibson assembly combining a EIF2A-Puro-T2A-GFP cassette replacing EIF2A:Puro in CROP-EIF2A-Puro(F+E)^38,68^. A BsmBI site within EIF2A::Puro-T2A-GFP was then deleted by site-directed mutagenesis, and the gRNA scaffold was modified to be compatible with direct-capture scPerturb-seq from 10x Genomics by inserting the capture sequence 1 within the stem loop of the gRNA scaffold^69^.

### Gene selection for perturbations

Genes to be perturbed in the CRISPR-Cas9 screen were selected from literature and data analysis of the scRNA-seq datasets (total n = 400). Genes included: (i) Top expressed TFs per cell type (ii) Literature-curated genes from GWAS on CHD and myopathies, as well as, heart development^70–73^; genes from Causal Variant to Gene Mapping Statistics (COGS)-analysis linking GWAS and HiC data; genes with implication in both heart disease and neurodevelopmental disorders^36,74–76^ (iii) Co-expression network analysis on the scRNA-seq atlas, using highly variable genes: top 5 genes by degree (undirected graph), betweenness, and eigenvalue centrality (iv) Positive control pluripotency genes (Supplementary Table 6). Except for TFs – which are often lowly expressed – genes to be perturbed were only included if they were expressed in more than 10% of cells with a minimum normalised and log-transformed expression level bigger or equal to 0.1 within cardiac progenitors, cardiomyocytes, fibroblast, endothelial cells or mesoderm in the scRNA-seq dataset. Furthermore, genes to be perturbed were filtered for essentiality^77^.

### gRNA design and library cloning

All gRNAs for perturbations were obtained from the Human CRISPR KO Library (GeCKO v2)^78^. Four gRNAs were selected for each gene. As negative control (NC) gRNAs, 16 non-targeting control (NTC) gRNAs were selected, as well as, 16 control gRNAs designed to target the CLYBL locus, serving as cutting negative control gRNAs (NTC_CLYBL). Oligonucleotide pools for gRNA libraries were checked for the presence of BbsI and replaced with alternative oligos if necessary. gRNA oligonucleotides consisted of spacer sequences and overhangs for amplification that included BbsI sequences compatible with Golden Gate cloning. Oligonucleotide pools were ordered from TWIST (Supplementary Table 18). Oligonucleotides were amplified (eight cycles) with compatible primers. The purified PCR product was cloned into the respective CROP-seq screening vector described above using BbsI and Golden Gate cloning. Electrocompetent bacteria (Lucigen, 60242-1) were electroporated (10LµF, 600LΩ, 1,800LV, EL=L184LVLcm^−1^) with purified ligation product and grown in a pool for 10Lh at 30L°C before extracting plasmid DNA (endotoxin free).

### Virus production and infection of human iPSCs

Lentiviruses were produced in HEK293T (ATCC, CRL-3216) cells grown in DMEM supplemented with 10% FBS, 1× GlutaMAX (Gibco, 35050061), 100LULml−1 penicillin–streptomycin (Gibco, 15140122) and 1× MEM nonessential amino acids (Gibco, 11140050) and coated using VSV-G. The day before transfection, HEK293T cells were plated at 80% confluency, plasmids were lipofected using Lipofectamine 3000 Transfection reagent (L3000001), and the cells were split 1:10 5Lh after lipofection. The supernatant was collected 3Ldays after lipofection, cell debris was pelleted at 200 g for 5Lmin at 4L°C, the remaining supernatant was spun at 28,000 g for 5Lh, and the virus pellet was resuspended in the appropriate volume of E8L+LRI. On the day of infection, human Cas9-expressing iPSCs were split 1:2.5 2Lh before infection using Accutase. Infections were performed overnight in E8L+LRI. Medium was replaced the next day with E8. EGFP positive iPSCs were sorted by flow cytometry to ensure >1,000x coverage of gRNA representation within sorted iPSCs, plated and recovered for a week before setting up cardioid differentiations.

### snMultiome-seq data generation and pre-processing

Cardioid nuclei from different time points were processed using the Chromium Next GEM Single Cell Multiome ATAC + Gene Expression Kit (10x Genomics - PN-1000285) according to the user guide (CG000338 Rev E). Two independent differentiations experiments of cardioids were grown with complementing time points to get the time-resolved data. Differentiation batch one was harvested at days 2, 6, 8, and 15; batch two at days 3, 4, 5, and 7. Multiome libraries (RNA and ATAC) were sequenced separately. Raw FASTQ files were processed using Cell Ranger ARC (v2.0.2; 10x Genomics) with default parameters. snATAC-seq data were processed using SnapATAC2^79^. Nuclei with <1,000 fragments or TSS enrichment score <4 were excluded. Gene activity matrices were generated by counting fragments overlapping promoter regions and gene bodies. The corresponding snRNA-seq data were processed using scanpy (v.1.11.5)^80^. Comparison of the two differentiation experiments revealed differences in differentiation progression based on pseudotime inferred using Palantir^43^: Cells collected on day 6 (Differentiation batch 1) exhibited higher pseudotime values than cells collected on day 7 (Differentiation batch 2), indicating that chronological sampling time was not directly equivalent between the two differentiation experiments. We therefore did not regress out differentiation batch, to avoid removing biologically meaningful variation associated with differences in differentiation progression. TF motif activity was inferred from snMultiome-seq data using pychromVAR^81^ (v.0.0.4) with motifs from the JASPAR 2024 database^82^. Bias-corrected motif accessibility deviation scores were calculated across cells.

### scRNA-seq and scPerturb-seq data generation and pre-processing

One cardioid differentiation was performed for unperturbed scRNA-seq, whereas scPerturb-seq data were generated from two independent cardioid differentiations initiated two weeks apart. Cells were dissociated at the indicated time points and fixed as described above. For scRNA-seq, fixed cells were processed with the ScaleBio scRNA-seq Kit (Beta v2) according to the manufacturer’s instructions (09Jan23). For scPerturb-seq, fixed cells were processed with the Cas9 CRISPR Guide Enrichment Kit v1.1 (ScaleBio - 955097) and the CRISPR Guide Enrichment Extended Throughput Kit v1.1 (ScaleBio - 945058) according to the manufacturer’s instructions (RevB 28NOV2023). Gene-expression and CROP-seq libraries for scPerturb-seq were generated separately, pooled at a 10:1 ratio and sequenced together. Cells collected on days 2–8 were processed across three reverse-transcription plates, each with three extension plates. Libraries from day 2–4 and 6 were sequenced together across three NovaSeq 6000 S4 runs with XP mode totaling 12 lanes (v1.5 - 200 cycles kit) with R1: 34, R2: 100, i5: 10 and i7:10 reads. Libraries from day 5 and 8 were sequenced across four NovaSeq 6000 S4 runs totaling 16 lanes (v1.5 - 200 cycles kit) with R1: 34, R2: 100, i5: 10 and i7:10 reads. Raw FASTQ files were processed using the ScaleRNA (v2.1.0; Scale Biosciences) and ScaleCRISPR (v1.0.0; Scale Biosciences) pipelines with default parameters (https://github.com/ScaleBio/ScaleRna, https://github.com/ScaleBio/ScaleCRISPR). For the day 2 time point only one cardioid differentiation was recovered after sequencing. Reads were aligned using STARsolo (v.2.7.11a), retaining only uniquely mapping reads (--soloMultiMappers Unique) for comparability with 10x Multiome data.

### gRNA quality control and assignment

Guides were assigned to cells with read purity ≥ 0.9, ≥ 100 reads, and ≥ 3 UMIs. Read purity was defined as the proportion of reads from the most abundant guide. Cells with multiple or no detectable gRNA assignments were excluded.

### RNA-seq normalisation, dimensionality reduction, integration, and cell type annotation

Pre-filtered scRNA count matrices were subjected to a unified quality control and preprocessing pipeline, with thresholds adjusted for sequencing depth across datasets (Supplementary Table 2). Doublets were removed using scDblFinder (v.1.16.0)^83^ and cell cycle scores were calculated using Scanpy score_genes_cell_cycle() with genes from Tirosh I. *et al*.^84^ Following mRNA quality control, and for scPerturb-seq, guide assignment, raw counts from snMultiome, scRNA-seq and scPerturb-seq datasets were merged. The top 4,000 highly variable genes were identified using (flavor=’seurat_v3’)^80,85^. Counts were normalised to the median library size of the combined dataset, log1p-transformed and subjected to PCA using highly variable genes. PCA embeddings were integrated across datasets using Harmony (v.1.2.4)^86^ (lamb=2, theta=2.5). A nearest-neighbour graph was constructed from the integrated space using 10 neighbours and 20 principal components. Cell types were manually annotated across the scRNA-seq, snMultiome-seq and scPerturb-seq datasets based on the integrated embeddings and expression of curated marker genes (Supplementary Table 4). Leiden clusters with mixed cell-type marker expression or low marker expression, indicative of doublets or empty droplets, respectively, were excluded.

### Trajectory inference with Palantir and CellRank2

Cell fate trajectories were inferred separately for each sequencing dataset using Palantir (v.1.4.1)^43^. Prior to trajectory inference, the gene set was restricted to highly variable genes, together with any genes annotated as perturbations and a curated list of congenital heart disease (CHD)-associated genes^27^, to focus the analysis on biologically relevant features. Diffusion maps were computed on the top 20 PCs using palantir.utils.run_diffusion_maps(), and a multiscale diffusion space was determined from the top 20 diffusion components (determine_multiscale_space()). A start cell was defined by randomly sampling from the Mesendoderm cells at day 2, and terminal states were similarly defined by randomly sampling one representative cell per specified terminal cell type (Cardiomyocytes, Fibroblasts, Endothelial cells, Ectoderm, Endoderm) at day 8 (scPerturb-seq) or 15 (snMultiome, scRNA-seq).

Palantir was then run (palantir.core.run_palantir()) using the selected start cell, terminal states, and a specified number of waypoints (3,000 for scPerturb-seq, 2,000 for snMultiome and scRNA-seq) to estimate pseudotime, terminal-state differentiation probabilities, and entropy for each cell. To assess cell fate probabilities with CellRank2, a 20-nearest-neighbour graph was constructed from 30 Harmony-corrected principal components. CellRank’s PseudotimeKernel used Palantir pseudotime to compute a transition matrix, and GPCCA identified six macrostates using 30 Schur components. Cardiomyocytes, fibroblasts, endothelial cells, ectoderm, and endoderm were specified as terminal states, and CellRank fate probabilities were calculated for each cell.

### Cell type preferential gene expression of congenital heart disease genes

For each gene, mean expression was computed across all cells of each cell type, then z-scored across cell types to quantify expression specificity. For each cell type, genes with a specificity z-score > 1 were ranked by mean expression, and the top 300 were retained as that cell type’s preferential gene set, favouring genes both enriched in and highly expressed by that cell type. A background gene universe was defined as all genes with mean log1p-transformed expression > 0.001 across the dataset. CHD-associated gene sets were obtained from Ma, X. R. *et al*.^27^, which assigned candidate genes to individual CHD traits, a pooled all-CHD category, and a *de novo* coding variant set (PCGC *de novo* variants)^36^. For each cell type-trait combination, we tested over-representation of the trait gene set within the cell type’s preferential gene set (vs. background) using a one-sided Fisher’s exact test (odds ratio > 1), with the 2×2 table built from trait/non-trait gene counts in the top-300 set and in the background. *P* values were Benjamini-Hochberg-corrected within each trait across cell types (FDR < 0.05 for significance; Supplementary Table 5).

### Quantification of perturbation effects on cell type abundance

For each perturbed gene and cell type, we tested whether cells carrying a targeting guide were over-or under-represented in that cell type relative to cells carrying NC guides. Guides targeting the same gene were pooled, as were all NC guides. For each perturbed gene-cell type pair, cells were cross-classified into a 2×2×B contingency table (cell type of interest vs. all others × perturbed gene vs. NC × cardioid differentiation batch, B strata) and tested for differentiation batch-stratified association using a two-sided Cochran-Mantel-Haenszel (CMH) test, which yields a common (Mantel-Haenszel) odds ratio and *P* value per pair. *P* values were corrected for multiple testing (Benjamini-Hochberg). For Fig. 2b, c we retained perturbed genes with FDR < 10LL and an absolute odds ratio (log2) ≥ 0.5 in at least two cell types.

### Clustering of cell type abundance effects

Perturbed genes were hierarchically clustered (SciPy; v.1.15.3) on their per-cell-type log2-odds-ratio (log-OR) profiles from the CMH tests above, using the same FDR < 10LL, odds ratio (log2) ≥ 0.5 in at least two cell types filter. Odds ratios of 0 occurred only for the under-represented cell type “Differentiation-impaired”. This cell type was excluded due to small cell numbers and resulting inflated log-Ors. Log-ORs were capped at 4 to limit the influence of near-complete-separation effects. For clustering only (not for display), cell types were collapsed into eight developmentally related groups – Mesendoderm; Cardiac mesoderm; Cardiac progenitors; Cardiomyocytes (early and mature); Fibroblasts (early and mature); Endothelial; Ectoderm (early, SOX2+, and mature); Endoderm (early and mature) – by averaging log-ORs across each group’s constituent cell types per target. The resulting matrix was clustered using Euclidean distance and Ward’s linkage, cutting the dendrogram at k = 7. Two clusters were further subdivided by re-clustering (Euclidean/Ward) their member targets alone, yielding two subclusters each (I/H and D/E).

### Differential gene expression analysis

Raw mRNA counts were aggregated across cells sharing the same gRNA, cell type, and cardioid differentiation batch to generate pseudobulk profiles. Low-quality pseudobulk samples were filtered in three steps: (1) pseudobulk samples with fewer than five cells per guide were excluded; (2) within each cell type, pseudobulk samples below the 0.5th percentile of total counts were removed as low-depth outliers, and potential *trans-*genes with fewer than 10 counts were excluded; (3) for robust downstream differential expression, perturbations were retained only if supported by at least three pseudobulk replicates per cell type. 389 perturbations were included in differential expression testing. Differentiation-impaired cells showed too little coverage to meet the above-described thresholds to include any perturbations for differential expression testing. Highly variable genes (HVGs) were identified per cell type using the seurat_v3 method^85^ in Scanpy on raw counts, selecting the top 10,000 HVGs per cell type. All gRNA-targeted genes were retained regardless of variability. Following Zhu *et al*.^87^, perturbed genes were randomly shuffled (fixed seed) and partitioned into consecutive chunks of 50 (final chunk holding the remainder) to facilitate batched downstream analysis. A binary target-by-chunk membership matrix was generated per cell type.

DESeq2^46^ (python implementation PyDESeq2 v.0.5.4^88^) was run independently per chunk on pseudobulk counts, comparing each perturbation target to NC guides (design formula: ∼ log10_n_cells + differentiation_batch + target) within the same cell type as:

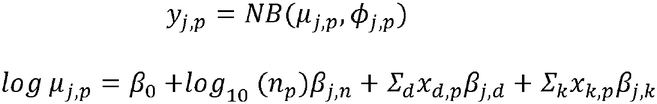

where *β*_0_ is the intercept, *j* indexes downstream genes and *p* indexes pseudobulk samples (a given perturbation in a given cell type). *n*_p_ is the number of cells in a pseudobulk *p,* with effect *β*_j,n_ on mean expression of gene *j. β*_j,d_ models the effect of differentiation batch (*x*_d,p_: indicator variable of differentiation batch of pseudobulk *p*). The effect of the perturbation *k* on mean expression of gene *j* is modelled by *β*_j,k_, where *x*_k,p_ is an indicator variable denoting the assignment of pseudobulk *p* to a perturbed gene *k*.

For each perturbed gene *k*, log2 fold-changes (*β*_j,k_) were estimated and tested against NC pseudobulks with a Wald test, fitting models independently per cell type to obtain context-specific estimates. To account for differing cell recovery across replicates, we report the z-score of the DE log-fold change (Wald statistic) unless otherwise noted, following Zhu *et al*.^87^:

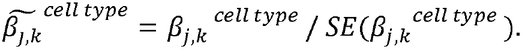

To assess *P* value calibration under the null, we performed resampling using only NC pseudobulks. In each of 10 iterations per cell type, 4 NC guides were randomly sampled without replacement (matching the 4 gRNAs per perturbed gene), with remaining NC samples as the control group; each comparison was processed through the same pseudobulk DESeq2 pipeline, retaining per-gene *P* values across iterations. For each cell type, the resulting null *P* value distribution (“null negative controls”) was compared via Q-Q plots (observed vs. expected −log10(p) under a uniform null) against 30 randomly sampled true targeting perturbations.

### Cell-type specificity and timing of abundance effects

Analysis was restricted to perturbed genes retained by the cell type abundance clustering analysis above. For each target, we identified the most upstream cell type – separated by branch – with a qualifying effect, excluding downstream occurrences inherited from an ancestor branch.

### Onset and persistence of *trans*-effects

For each perturbation-branch pair, we defined the onset as the first cell state along the trajectory with ≥10 significant genes and recorded its trajectory position and number of significant genes. Persistence was assessed by scanning from the onset to the terminal state and recording the most differentiated state that met the same significance threshold, irrespective of significance in intervening states. Perturbations with no significant downstream state were classified as “only here”; otherwise, they were assigned the furthest-affected state. For perturbations with onset in near-terminal or terminal cardiac fates (cardiomyocytes, fibroblasts or endothelial cells), we quantified the lineage breadth of *trans*-effects. Perturbations were grouped by onset lineage, and the proportion affecting one, or at least two cardiac lineages was calculated.

### Quantification of total *trans*-effects and perturbation target gene expression along pseudotime

For each perturbed gene, differentially expressed downstream genes were identified from pseudobulk DESeq2 results (adjusted P < 0.05, |log2 fold-change| ≥ 1) computed per perturbation and cell type. To increase statistical power for temporal analysis, cell types were aggregated into coarse lineage bins and significant downstream genes were pooled across all cell types within a lineage bin. Target-lineage combinations with fewer than 10 significant downstream genes were excluded. For each target *t* and cell type *i* we defined a gene set *G*_i,t_ based on cell type-level DE testing as:

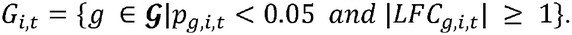

Cells were assigned to coarse lineage bins based on their cell type annotation. Within each lineage, the total number of pseudotime bins (n = 40 across the dataset) was allocated proportionally to the number of cells in that lineage, with fractional remainders distributed using a largest-remainder method to ensure the bin total was preserved. Cells within each lineage were then ranked by Palantir pseudotime and divided into equally sized, sequentially ordered bins.

Raw UMI counts were summed per pseudotime bin and per pseudobulk group, defined by the combination of perturbation target and lineage bin. In parallel, the percentage of cells expressing each gene (count > 0) was computed per group. Pseudobulk groups represented by fewer than 5 cells were discarded. Targets were retained if they were represented in at least 25 pseudotime bins. Pseudobulk counts were normalised by total counts per group.

For each perturbation target and lineage bin, pseudotime bins containing both perturbed cells and NC cells were matched, and a log2 fold-change was computed per downstream gene as log2(normalised target expression + 1) − log2(normalised NC expression + 1). To summarize the overall transcriptional impact of a perturbation independent of the direction of individual genes, we defined a total effect size per target, lineage bin, and pseudotime bin as the root-sum-of-squares of the downstream gene log2 fold-changes (Supplementary Table 9):

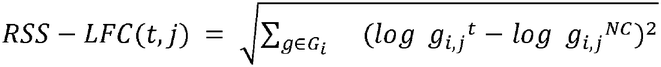

with pseudotime bin *j*, cell type *i* and perturbation *t*. *g_i,j_^t^* and *g_i,j_^NC^* denote normalised gene expression levels of gene *g* in pseudotime bin *j* and cell type *i* with perturbation of gene *t* or NC, respectively.

To compare dynamics across perturbations with different expression and effect magnitudes, log1p-transformed target expression and RSS log2 fold-change were independently normalised by their respective maxima across all pseudotime bins and lineages for each target. Targets were retained if they had a nonzero RSS log2 fold-change in at least one bin. Profiles were required to contain at least one bin with ≥10 genes contributing to the RSS and at least one bin with an absolute scaled RSS log2 fold-change ≥0.3. We further excluded profiles with insufficient target expression accompanying early perturbation effects. Separately for the pluripotent and cardiac progenitor stages, we considered bins with unscaled RSS log2 fold-change ≥0.3 and available target expression measurements, and calculated the fraction with log1p-transformed target expression ≤0.3. Profiles were excluded if this fraction exceeded 50% in either stage. Stages without evaluable bins did not cause exclusion.

### Clustering of response-timing along pseudotime

Cardiomyocyte and fibroblast lineage profiles were analysed jointly, with each perturbation represented separately in each lineage. Each profile was encoded as a concatenated vector of normalised target expression and RSS log2 fold-change across ordered pseudotime bins spanning pluripotent, cardiac progenitor, and terminal stages. To align terminal-stage profiles, the eight fibroblast bins were mapped to evenly spaced positions, rounded to integer indices, along the 15-bin cardiomyocyte axis. Unobserved positions were retained as missing values. Features were mean-centred and scaled to unit variance, and probabilistic principal component analysis (PPCA) was used to accommodate missing observations and obtain five components. Hierarchical clustering was performed on Euclidean distances between the five-component PPCA scores using Ward’s minimum-variance method, and the dendrogram was cut into six clusters. For visualization, the same PPCA scores were embedded in two dimensions using UMAP (Euclidean distance, 12 neighbours, minimum distance 0.1).

### Training scDORI on snMultiome data

Enhancer-mediated GRNs were inferred jointly from snRNA-seq and snATAC-seq data using scDORI (v1.0.0). Analyses were restricted to 7,155 genes for computational scalability. Genes included 666 TFs drawn from 3,000 highly variable genes, 300 highly variable TFs, 400 perturbed genes, genes differentially expressed following target-gene knockout, and cardiac disease genes. Genes were retained if mean expression exceeded 0.2, and TFs if mean expression exceeded 0.1, in at least one cell type.

TF-binding motifs from HOCOMOCO v14^89^, JASPAR 2026 CORE^90^ and CIS-BP^91^ were mapped to accessible peaks using the tangermeme (v.0.4.3) implementation of FIMO^92^ (significance threshold 0.001). Motif-peak mappings were used to calculate *in silico* ChIP-seq scores for activators and repressors following the scDORI preprocessing workflow.

The scDORI topic model was fitted with 40 topics and a batch size of 16. In phase one, the encoder, topic-peak decoder, peak-gene links and topic-TF decoder were trained for up to 1,000 epochs (five warm-up epochs, early stopping after 25 epochs without validation-loss improvement; L1 = 10LL for the topic-peak decoder, other parameters default). In phase two, topic-specific scDORI-inferred GRNs were reconstructed from the fitted topics and used to reconstruct RNA expression, for up to 1,000 epochs (early stopping after 10 epochs without improvement).

### Construction of the consensus GRN combining snMultiome and scPerturb-seq

This network was constructed independently of scDORI’s topic-specific GRN, using only scDORI’s peak-gene linkage matrix and *in silico* ChIP-seq scores together with scPerturb-seq evidence. First, cell-type-specific GRNs were constructed from three multiome-derived components: (i) signed *in silico* ChIP-seq scores from scDORI, obtained by summing activator (positive) and repressor (negative) *in silico* binding scores for each TF-peak pair, with near-zero values rounded to zero using NumPy (v.1.26.4) (isclose() function) (ii) scDORI peak-gene linkage scores, and (iii) mean chromatin accessibility for each peak and cell type, calculated from snATAC-seq.

Second, these multiome-derived GRNs were intersected with scPerturb-seq trans-effects. scPerturb-seq differential expression results were restricted to genes with FDR < 0.1, and to TFs and genes represented in the scDORI model. For each cell type and TF with significant trans-effects in scPerturb-seq, we identified peaks bound by the TF (nonzero *in silico* ChIP-seq score) and linked these peaks to differentially expressed genes using the scDORI peak-gene linkage matrix. A gene was considered a candidate TF target if it was linked to at least one TF-bound peak.

Third, for each remaining TF-gene pair, we calculated a multiome-supported edge weight as the mean, across linking peaks, of the product of three terms: (i) cell-type-specific peak accessibility, (ii) scDORI peak-gene linkage score, and (iii) the TF’s *in silico* ChIP-seq binding score:

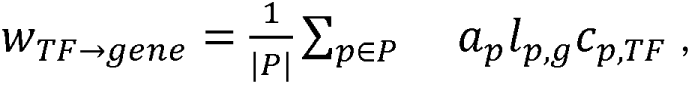

where *a_p_* is the mean chromatin accessibility of peak *p* in a given cell type, *l*_p,g_ is the scDORI peak-gene linkage score, *c*_p,TF_ is the signed *in silico* ChIP-seq score and *P* is the set of peaks with non-zero values for all three terms. TF-gene pairs with a resulting score of zero were discarded. For the retained edges, the final edge weight was set to the log fold change measured by scPerturb-seq differential expression, as this provides direct experimental evidence of the magnitude and direction of the TF perturbation effect. The resulting edges with non-zero weight constituted the consensus snMultiome-scPerturb-seq GRN.

### Classification of direct and indirect regulatory links based on TF-chromatin-binding evidence

To determine whether *trans*-effects identified by scPerturb-seq were supported by chromatin-binding evidence, each TF-gene effect was queried against the corresponding cell-type-specific consensus snMultiome-scPerturb-seq GRN using NetworkX (v.3.4.2). Links were classified as **direct** when the TF and target were connected by a single edge in the consensus GRN, **multi-hop** when connected by a directed path of two or more edges, and **indirect link - no overlap** when there was no path between TF and downstream gene in the consensus GRN. Direct links were further classified as sign-concordant or sign-discordant depending on whether the direction of effect from scPerturb-seq was the same as the direction of effect inferred.

### Permutation testing of multiome-derived GRN edges

To test whether the overlap between the scPerturb-seq-derived and multiome-derived TF-gene regulatory networks exceeded chance expectation, both networks were first restricted to the TFs and genes shared between them. Each scPerturb-seq TF-gene edge (FDR < 0.1) was then classified as a direct hit if the identical TF-gene edge was present in the multiome network, or as a multi-hop hit if the gene was reachable from the TF via a directed path of at most 10 edges in the multiome network, identified by bounded breadth-first search; edges satisfying neither criterion were classified as not found. To generate a null distribution for the number of direct and multi-hop (direct + multi-hop, “reachable”) hits expected by chance, we permuted the target-gene identity of each scPerturb-seq edge by sampling uniformly, with replacement, from the full set of shared genes, not preserving the out-degree. For each of 1,000 permutations, direct and reachable overlap with the (unpermuted) multiome network were recomputed. Empirical, one-sided *P* values were calculated as *P* = (1 + number of permutations with overlap ≥ observed)/(1 + number of permutations), and permutation z-scores were reported as (observed − null mean)/null standard deviation. This procedure was applied separately within each cell type.

### Pairwise GRN similarity

Pairwise similarity between cell-type GRNs was quantified separately for edges (TF-gene links) and nodes (TFs & genes) using the Jaccard index (intersection over union) and the overlap coefficient (intersection over the smaller set size), and visualised as combined heatmaps with the two metrics shown in upper/lower triangles. The Jaccard index was calculated as

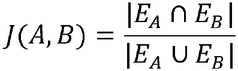

where *E*_A_ and *E*_B_ denote the edge/node sets of the GRNs for cell types *A* and *B*, respectively. Similarly, the overlap coefficient (Szymkiewicz–Simpson coefficient) was computed as:

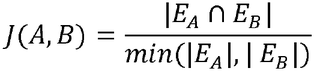

### TF-TF GRN rewiring across differentiation

Cell-type-specific consensus GRNs (TF-only subnetwork restricted to TF-TF edges) were compared across cell types and along two differentiation branches, corresponding to the cardiomyocyte and fibroblast lineages, to characterise regulatory rewiring during cardiac organoid differentiation. Pairwise metrics were computed between each pair of consecutive cell types along a branch (ct1→ct2). For both TFs (graph nodes) and TF-TF links (graph edges), we quantified network size, the number and fraction of elements lost (present in ct1 but absent in ct2) and gained (present in ct2 but absent in ct1), and the number and fraction of shared elements (present in ct1 and ct2). Metrics were summarised per transition as stacked bar plots of TF and TF-TF link turnover (gained, lost, and retained). This analysis was repeated using cell-type-specific GRNs inferred from snMultiome and using scPerturb-seq-based GRNs, to confirm that observed rewiring patterns were not driven by a single data modality.

### Cell-type specificity of TF out-degree

For each TF and cell type, row-wise z-scores of out-degree were computed across cell types to identify TFs with regulatory influence in specific cell types. TFs were hierarchically clustered based on these z-scores.

### Quantification of activating versus repressing TF-downstream gene regulatory bias

Cell-type-resolved GRNs connect each TF to its downstream genes, with edge sign (log fold change from DE testing) defining regulatory direction: negative for activating, positive for repressing. For each TF-cell type pair, we counted activating and repressing edges, classifying pairs as activating, repressing, or dual-role, and summarised bias as the fraction of activating edges (visualised as dot plots; colour, activating fraction; size, edge count; TFs clustered by average linkage, cell types ordered by developmental stage).

### Hierarchical clustering of pairwise correlation of perturbation trans-effects to identify co-regulator clusters

Perturbation-cell type conditions were filtered to require ≥10 significant (FDR < 0.1) *trans*-effect genes, retaining 213 perturbations. Only trans-effect genes significantly differentially expressed (FDR < 0.1) in at least one passing condition were retained. For each retained perturbation-cell type condition we used the DESeq2 Wald statistic (z-scored log-fold change) to build a conditions × genes matrix, with pairwise-complete observations handling genes not tested in a given condition. Pairwise Pearson correlations between conditions were hierarchically clustered by average linkage with Euclidean distance (scipy.cluster.hierarchy.linkage), and cut into k = 50 clusters (fcluster, maxclust criterion). Two of the resulting clusters (Mixed-8 and Mixed-11) captured perturbation-cell type conditions with only weakly correlated *trans*-effects and were excluded or used as negative controls in downstream analysis. The resulting co-regulator clusters contained 125 unique perturbations, with 88 unique perturbations falling exclusively into one of Mixed-8 and Mixed-11.

### Cell-fate, co-regulator and response-timing cluster overlap and enrichment analysis

Overlaps were calculated using unique perturbed genes assigned to cell-fate, co-regulator and response-timing clusters. Co-regulator cluster assignments to Mixed-8 and Mixed-11 were excluded because these clusters showed weakly correlated effects across cell types. For overall gene-set overlaps, response-timing cluster genes were pooled across cardiomyocyte and fibroblast lineages and counted once. Cluster-level comparisons retained lineage-specific response-timing cluster assignments, with overlap quantified as the fraction of genes in each row cluster shared with the corresponding column cluster. Heatmap rows and columns were ordered independently using complete-linkage hierarchical clustering.

Enrichment of CHD-associated genes was evaluated separately for perturbations with effects on cell-fate, perturbations with ≥ 10 *trans*-effects and perturbations in co-regulator clusters, using one-sided Fisher’s exact tests. The background comprised all 389 genes included in differential-expression testing. For each of four tests (CHD, Multiple, DCM_HCM, not disease), disease annotation frequencies within the gene set were compared with those among the remaining background perturbations. P values were adjusted across the four tests using the Benjamini–Hochberg procedure. Fold enrichment was calculated as the proportion of disease-associated genes within each set divided by the corresponding proportion in the full background.

### Recurrence of cardiac progenitor co-regulator pairs in differentiated cell types

To assess the conservation of co-regulator relationships across clusters and cell types, we performed a pairwise recurrence analysis of genes represented within the same cell type and co-regulator cluster. For each co-regulator cluster, we extracted the set of unique perturbed genes in cardiac progenitors and generated all possible unordered combinations of two distinct genes within cardiac progenitors. A gene pair was considered recurrent if both genes co-occurred in at least one additional cluster in a differentiated cardiac cell type (one of (early) Cardiomyocytes, (early) Fibroblasts, Endothelial cells), taking into consideration the associated cell types. For each pair, we recorded the corresponding cell-type annotations. The originating cluster was included in all recurrence calculations. To quantify the recurrence of cardiac progenitor co-regulators at the co-regulator cluster level we counted all regulators that had at least one recurring co-regulator in at least one differentiated cell type versus all regulators that had noe recurring co-regulators.

### Metaclustering into co-regulator cluster families

Co-regulator clusters comprising fewer than two distinct perturbed genes were excluded from subsequent co-regulator-family-level analyses. One cluster not satisfying this criterion (co-regulator cluster “Mes-Cp-Fib-1”) was manually retained. This cluster contained a single perturbation in three cell types, but was retained as we observed correlation already at the cardiac progenitor stage that could partially explain the later convergence in Cm-6. Pairwise similarity between co-regulator clusters was quantified in two ways: (i) the mean pairwise Pearson correlation between member perturbation profiles (diagonal excluded), and (ii) the Jaccard index between each co-regulator cluster’s set of unique perturbations. Co-regulator clusters were grouped into families by average-linkage hierarchical clustering (Euclidean distance) of the Jaccard similarity matrix, cut into *k* = 14 groups (scipy.cluster.hierarchy.fcluster(), maximum-clusters criterion). A co-regulator cluster family was kept for downstream analysis if the maximum pairwise Jaccard index among its constituent co-regulator clusters was ≥ 0.3, with two exceptions: C6, an example of loss, and C7, an example of propagation on closer inspection. Hierarchical relationships between co-regulator clusters and their descendant co-regulator clusters emerging later in differentiation were additionally curated to summarize the branching structure underlying cluster diversification within co-regulator cluster families.

### Loss of correlated *trans*-effects

Loss of correlated *trans*-effects in response to differentiation were identified as cardiac progenitor co-regulator clusters that exhibited low perturbation set overlap (Jaccard < 0.1) with all other co-regulator clusters, specifically indicating Cp-Fib-3. For each perturbed gene, the number of significant *trans*-effects (unique differentially expressed genes, adjusted P < 0.1) detected in cardiac progenitors was compared to the maximum number detected in any other cell type; perturbations with a ratio ≥ 2.5 were classified as progenitor-restricted, and those below this threshold as having effects sustained into later cell states, despite losing their co-regulatory relationships.

### Preranked gene set enrichment analysis

To characterise biological processes associated with each co-regulator cluster, gene set enrichment analysis was performed using aggregated transcriptional effects across its perturbation conditions. Downstream genes were included if significantly differentially expressed (FDR < 0.1) in at least one condition within the co-regulator cluster. For each gene, we calculated the fractions of conditions with positive and negative Wald statistics, normalised by the total number of conditions. We used the larger fraction (dominant direction of regulation) as a measure of regulatory consistency. Genes were ranked by the median Wald statistic across conditions, multiplied by this consistency score, thereby prioritising genes with strong and consistent effects across the co-regulator cluster.

Genes were ranked in descending order of the weighted score and submitted to preranked gene set enrichment analysis (GSEA) using GSEApy (v.1.1.13) (gp.prerank()). Enrichment was tested against KEGG pathway gene sets (KEGG_2021_Human) and GO terms (GO_Biological_Process_2025) retrieved from Enrichr, retaining gene sets with between 10 and 500 members. Enrichment significance was assessed using 1,000 permutations of the gene ranking, with a weighted enrichment statistic (weight = 1, equivalent to standard GSEA). Gene sets with a Benjamini-Hochberg FDR < 0.1 were considered significantly enriched. Enrichment was run independently for each co-regulator cluster.

Enriched terms were formatted for readability and manually curated to exclude a list of terms not relevant for cardioids (Supplementary Table 13). Terms associated with fewer than four overlapping genes were excluded from downstream summaries.

### Extracting GWAS-evidence genes from Open Targets

Disease ontology data were retrieved from the Open Targets Platform (GraphQL API; https://api.platform.opentargets.org/api/v4/graphql) using the search term “congenital heart disease”. The associated disease identifier (EFO ID) and name were extracted. Descendant disease terms were obtained by querying the ontology hierarchy for the selected root disease using its EFO ID. Additionally, we included dilated and hypertrophic cardiomyopathy, as well as negative control diseases (carotid artery disease, coronary artery disease, benign smooth muscle neoplasm, chronic kidney disease, celiac disease, deep vein thrombosis, age-related macular degeneration).

For the curated list of EFO and MONDO disease identifiers, we queried the associatedTargets field to obtain all reported target associations together with overall association scores and evidence-type-specific datatype scores. We retained only associations supported by genetic evidence, defined as Open Targets datatype categories genetic_association, gene_burden, or somatic, following Zhu *et al*^87^. For each target, the maximum score across these genetic evidence categories was recorded as the genetic evidence score. Associations were included if at least one genetic evidence datatype score was greater than or equal to 0.1. For each retained association, we extracted disease identifier, disease name, gene symbol, Ensembl target identifier, gene name, overall association score, genetic evidence score, and contributing genetic evidence datatypes.

### Preranked gene set enrichment analysis for disease genes

Genes were ranked in descending order of the weighted score and submitted to preranked gene set enrichment analysis (GSEA) using GSEApy (gp.prerank()). Enrichment was tested against each disease category (curated list from Ma *et al*^27,36^, OpenTargets^93^) retaining gene sets with between 10 and 500 members.

Enrichment significance was assessed using 1,000 permutations of the gene ranking, with a weighted enrichment statistic (weight = 1). Gene sets with a false discovery rate (FDR q-value) below 0.1 were considered significantly enriched. Enrichment was run independently for each co-regulator cluster. Terms associated with fewer than four overlapping genes were excluded from downstream summaries.

### Co-regulator cluster propagation trees

To relate co-regulator clusters arising at earlier stages of differentiation to those observed in more differentiated cell types, co-regulator clusters were manually organised into hierarchical “propagation trees,” each rooted at an early-stage co-regulator cluster (for example, a cardiac-progenitor co-regulator cluster) and connected through one or more branches to descendant co-regulator clusters representing divergent or shared partitioning of the root co-regulator cluster’s regulators at later cell states.

For every parent-child pair in a tree, we computed the mean pairwise Pearson correlation between *trans*-effects (z-score of log2 fold-changes) of constituent perturbations, in addition to each co-regulator cluster’s own within-cluster mean pairwise correlation. For each root co-regulator cluster, perturbed genes were further classified by their fate along the tree – present only in the root, restricted to one descendant branch, shared across multiple branches, or introduced in a descendant co-regulator cluster and therefore absent from the root – to visualize the conservation and diversification of individual regulators across differentiation.

Disease and KEGG pathway terms were required to pass a false-discovery-rate threshold (Benjamini-Hochberg FDR < 0.1); the most positively and most negatively enriched terms by normalised enrichment score (NES) were retained per cluster (up to 30 up-and 30 downregulated KEGG terms; up to 20 up-and 20 downregulated disease/phenotype terms). For the retained disease terms, leading-edge gene membership was summarised per cluster as a gene-by-term matrix indicating, for each gene, whether it belonged to the leading edge of a significantly up-or downregulated term (or neither). For KEGG pathways, we listed the enriched/depleted pathways, but not the leading-edge genes.

### Statistics and reproducibility

The “Differentiation-impaired” cell type was excluded from cell-fate clustering due to very small cell numbers. Cutoffs for defining high-quality cells were set as described above. Enrichment of disease genes among highly expressed and cell-type-specific genes was performed using a one-sided Fisher’s exact test. Differential gene expression testing in the single-cell data was performed using PyDESeq2. Differential abundance of cell types was assessed using two-sided Cochran-Mantel-Haenszel tests, stratified by cardioid differentiation batch, except for cardiac mesoderm, where only one batch was recovered. Multiple-testing correction was performed using the Benjamini–Hochberg method, and adjusted *P* values are reported as FDRs. Significance thresholds are reported for each analysis in the relevant methods section. Significance values are reported as: ns (not significant, adj. P ≥ 0.05), * (adj. P < 0.05), ** (adj. P < 0.01) or *** (adj. P < 0.001). All box plots shown in this study show the centre line as the median, box limits indicate 25th and 75th percentiles, and whiskers indicate 1.5× the interquartile range; all outliers are displayed.

## Data availability

Raw sequencing data and processed data will be uploaded to GEO. Accession codes will be made available upon publication. Data can be shared upon request.

## Code availability

All custom scripts and pipelines used for data processing and analysis will be made available upon publication. Code can be shared upon request.

## Acknowledgements

We thank all members of the groups of L.M.S., O.S. and B.C. for feedback and technical expertise. We thank Stefan Schrod, Martin Rohbeck, Jana Braunger, Britta Velten, Jan Gleixner, Manu Saraswat and Philipp Schäfer for discussions related to data analysis. We thank the Genomics Core Facility (GeneCore) at EMBL, the Next Generation Sequencing core facility at DKFZ and the NGS Competence Center Tübingen (NCCT) for consultation and sequencing. We thank the Flow Cytometry Core Facility at EMBL for support and flow cytometry services.

## Author contributions

D.L., K.M., B.C., O.S. and L.M.S. designed the study, analysed data and wrote the manuscript with input from all authors. D.L. and H.J. performed experiments with support from M.K.. K.M. and S.M. performed data analysis, with support from D.L.. K.U. performed data analysis for pilot screening experiments. K.U., L.R.-G., A.M.-V. and J.H. provided input for perturbation gene selection. D.S. provided scientific guidance. B.M., M.M., W.H., A.P., J.P.R., B.C., O.S. and L.M.S. acquired funding, contributed to data interpretation and provided scientific guidance and supervision.

---

**Extended Data Figure 1.**
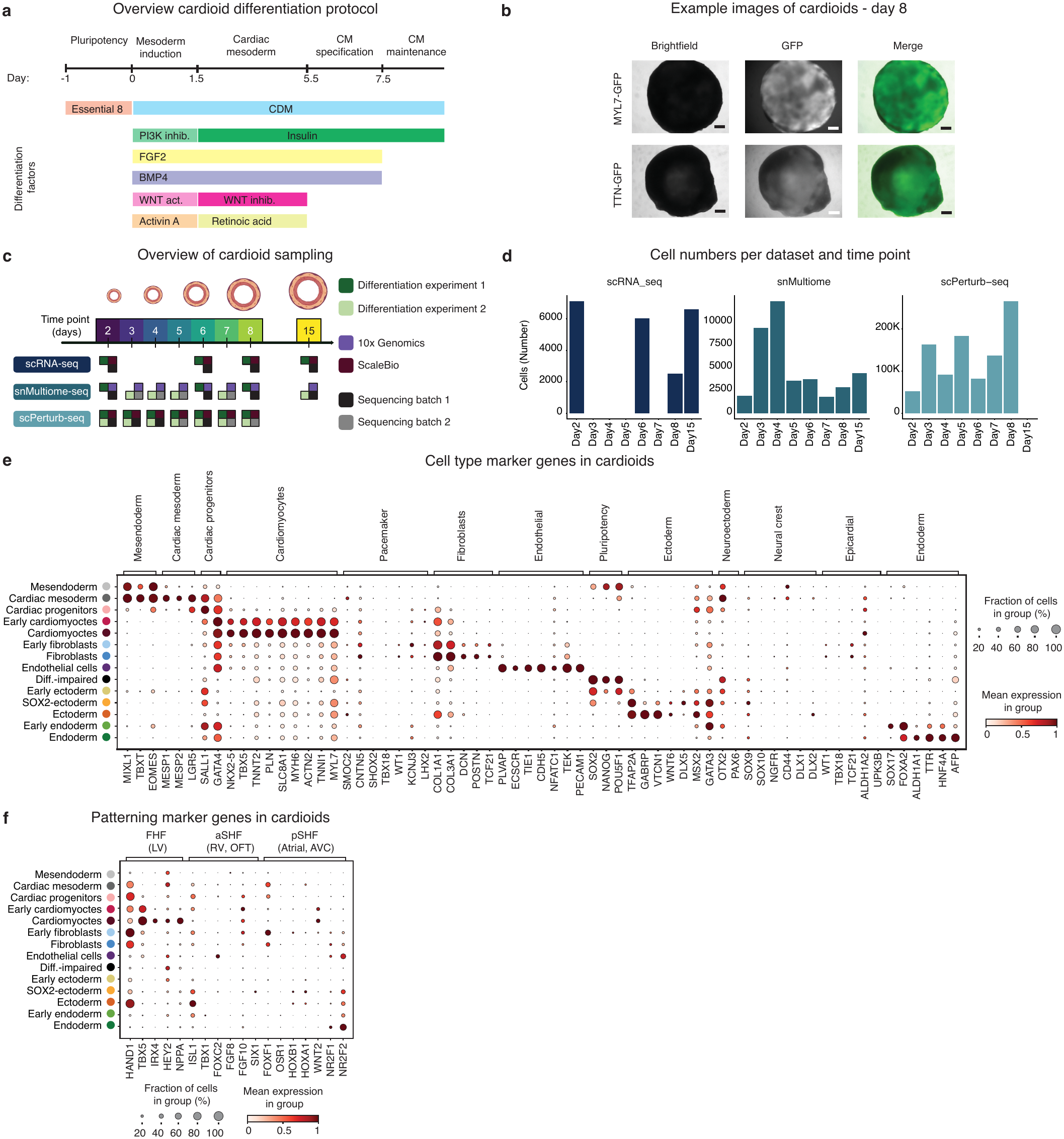
Cardioid differentiation protocol, time-resolved sampling details and marker gene expression. a, Overview of cardioid differentiation protocol. **b,** Example images of cardioids at day 8. iPSCs with endogenously tagged MYL7-GFP or TTN-GFP were used to set up cardioids, reporting on cardiomyocyte maturation. Scale bar = 200 µm. **c,** Overview of cardioid sampling. Number of differentiation experiments, single cell platforms and sequencing runs are indicated. **d,** Number of cells per dataset and time point. **e, f,** Dotplot of cell type marker gene expression **(e)** and patterning marker genes **(f)** across the annotated cell types. Scaled expression (colour) and percentage of cells expressing a marker (size) are shown.

**Extended Data Figure 2.**
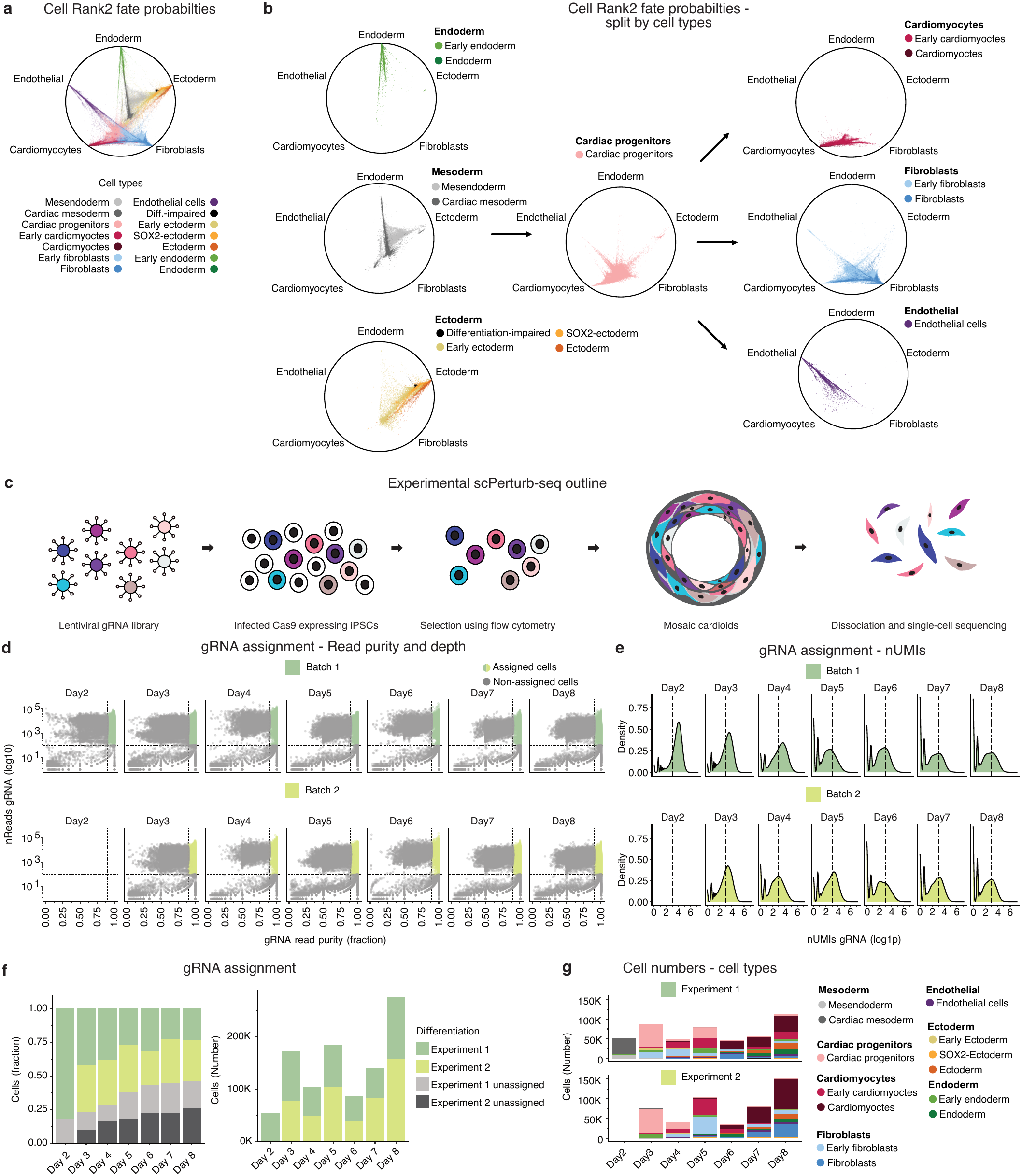
Fate predictions and quality control of scPerturb-seq dataset. a, Fate probabilities calculated with CellRank2 with colour denoting cell cycle. **b,** Fate probabilities calculated with CellRank2 split by cell type. **c,** Experimental workflow for the scPerturb-seq dataset. **d, e,** gRNA assignment based on RNA read depth (≥100 reads) and purity (≥90%) **(d)** and number of UMIs (≥3) **(e)** of the detected CROP-seq vector construct. **f,** Fraction of cells (left) and cell counts (right) of cells with assigned gRNAs in the scPerturb-seq dataset. **g,** Annotated cell types for cells assigned with a gRNA in both independent differentiation experiments in the scPerturb-seq dataset.

**Extended Data Figure 3.**
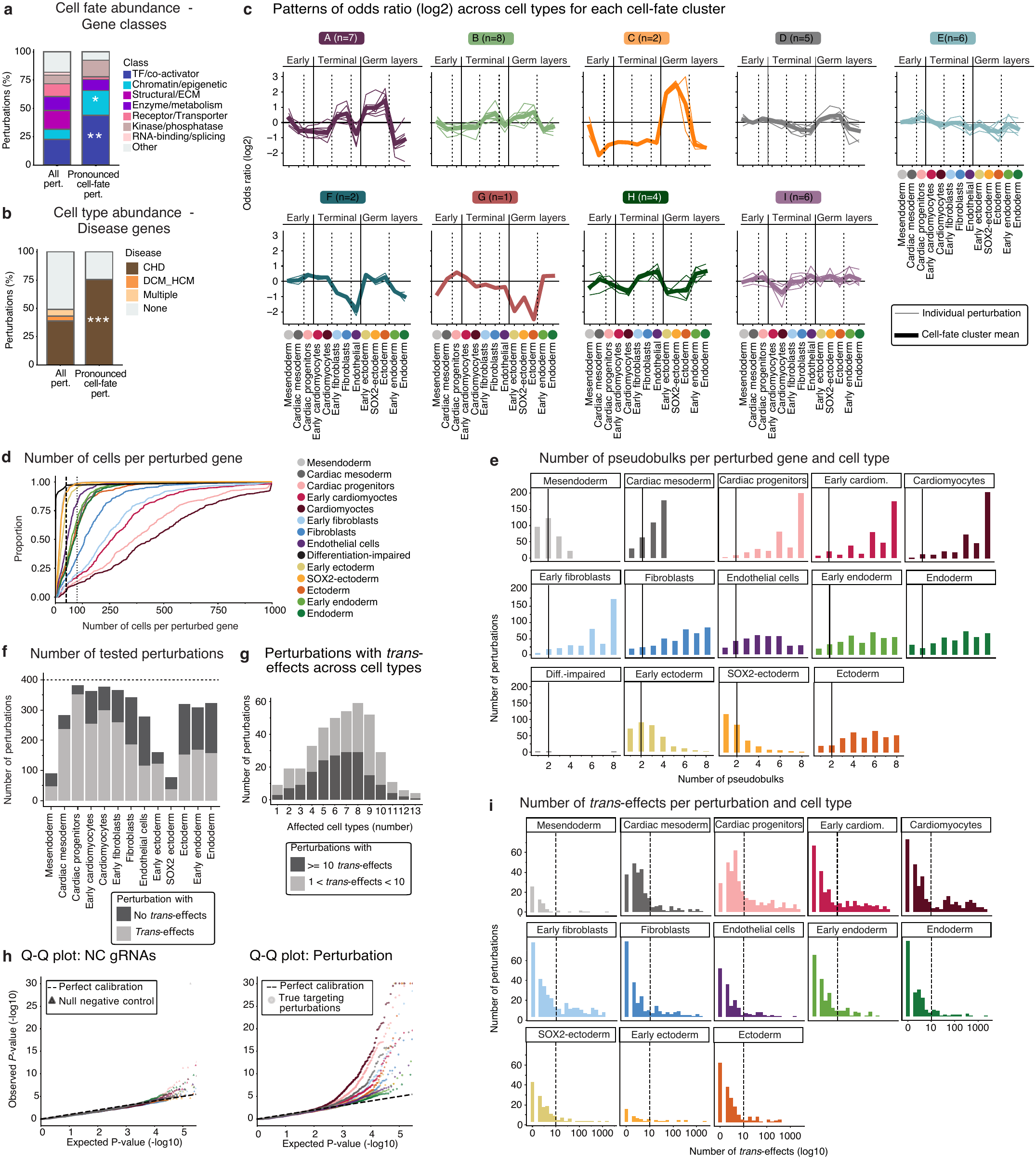
Cell fate abundance changes and differential gene expression analysis with DESeq2 to identify *trans*-effects. a-b, Relative numbers of class annotations **(a)** and disease annotations **(b)** of pronounced cell-fate abundance changing genes. One-sided Fisher’s exact test for enrichment. (TF \*\**P* = 0.00962; Chromatin/epigenetic \**P* = 0.01978; CHD \*\*\**P* = 2.27 x 10^-6^). **c,** Patterns of odds ratios (log2) across cell types within individual cell-fate clusters. Coloured thin lines indicate patterns per perturbation. Coloured bold lines indicate the mean pattern per cluster. Dotted lines indicate cell type and branch transitions. **d,** Cumulative distribution of number of cells per perturbation stratified by cell types (colour). 50 and 100 cells per perturbed gene are highlighted by vertical dotted lines. **e,** Number of pseudobulk replicates per perturbation and cell type used for DESeq2 testing (x-axis, replicates; y-axis, number of perturbed genes). Pseudobulk replicates were generated per gRNA (n = 4) and independent differentiation experiment (n = 2), yielding up to eight pseudobulk replicates per perturbation. Perturbations that had ≥3 pseudobulk replicates in a cell type were included in the testing. Low-depth outliers (bottom 0.5% of total counts) and potential *trans*-genes with <10 counts were excluded; ≥5 cells were required per pseudobulk replicate. **f,** Number of perturbations included in DE testing with and without *trans*-effects per cell type. **g,** Number of cell types affected by each perturbed gene, split by more or fewer than ten *trans*-effects. **h,** Quantile-quantile plots comparing observed versus expected *P* values (-log10) for null and true differential expression tests in each cell type. Dot colour indicates cell type, same as in **(d)**. The dashed line indicates perfect calibration (expected uniform distribution under the null hypothesis). Left: triangles represent null negative control tests, 10 random tests, four randomly sampled NC gRNAs. Right: points represent true targeting perturbations, 30 random tests shown. **i,** Histogram of number of *trans*-effects (log10) for perturbed genes. At least 10 *trans*-effects were required to include a perturbed gene in downstream analysis.

**Extended Data Figure 4.**
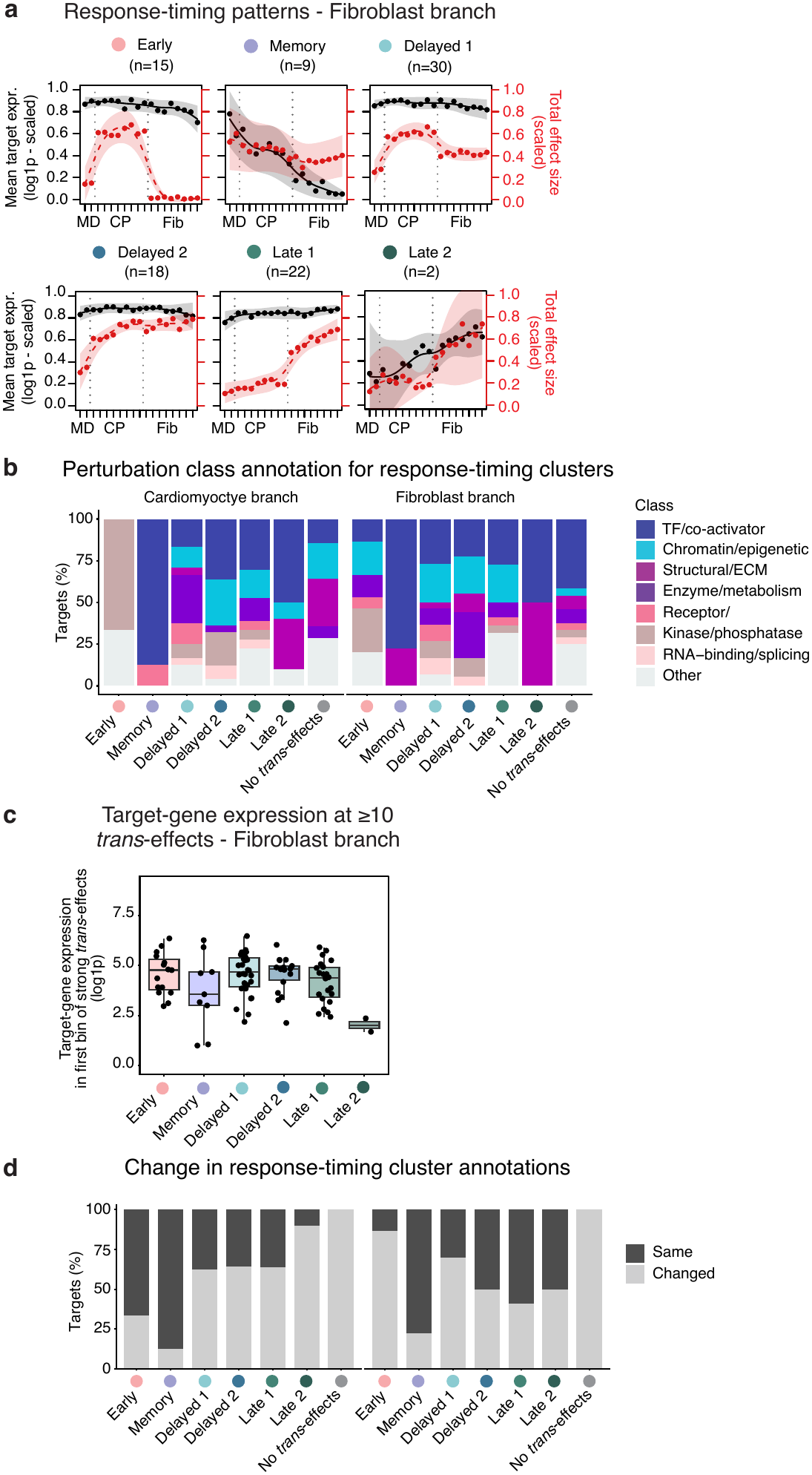
Response-timing cluster dynamics in the fibroblast branch, and comparison of response-timing cluster assignment between branches. **a**, Response-timing patterns of the clusters identified shown for the fibroblast branch. For each response-timing cluster, mean max-scaled target-gene expression (log1p; black, left axis) and mean max-scaled total effect size (root sum of squared log2 fold changes; red, right axis) across member perturbations are shown per pseudotime bin, from mesoderm (MD) through cardiac progenitors (CP) to fibroblasts (Fib). Points: bin means; lines: spline fits with shaded 95% confidence intervals; n: number of perturbations per cluster. **b,** Stacked bar plots showing the percentage of perturbations (y-axis) belonging to each functional gene class within each response-timing cluster, shown separately for the cardiomyocyte branch (left) and fibroblast branch (right). **c,** Target-gene expression at the onset of strong *trans*-effects (≥10) per response-timing cluster in the fibroblast branch. **d**, Stacked bar plots showing the percentage of perturbations (y-axis) whose response-timing cluster annotation remained the same or changed between the two branches, shown for each response-timing cluster in the cardiomyocyte branch (left) and fibroblast branch (right).

**Extended Data Figure 5.**
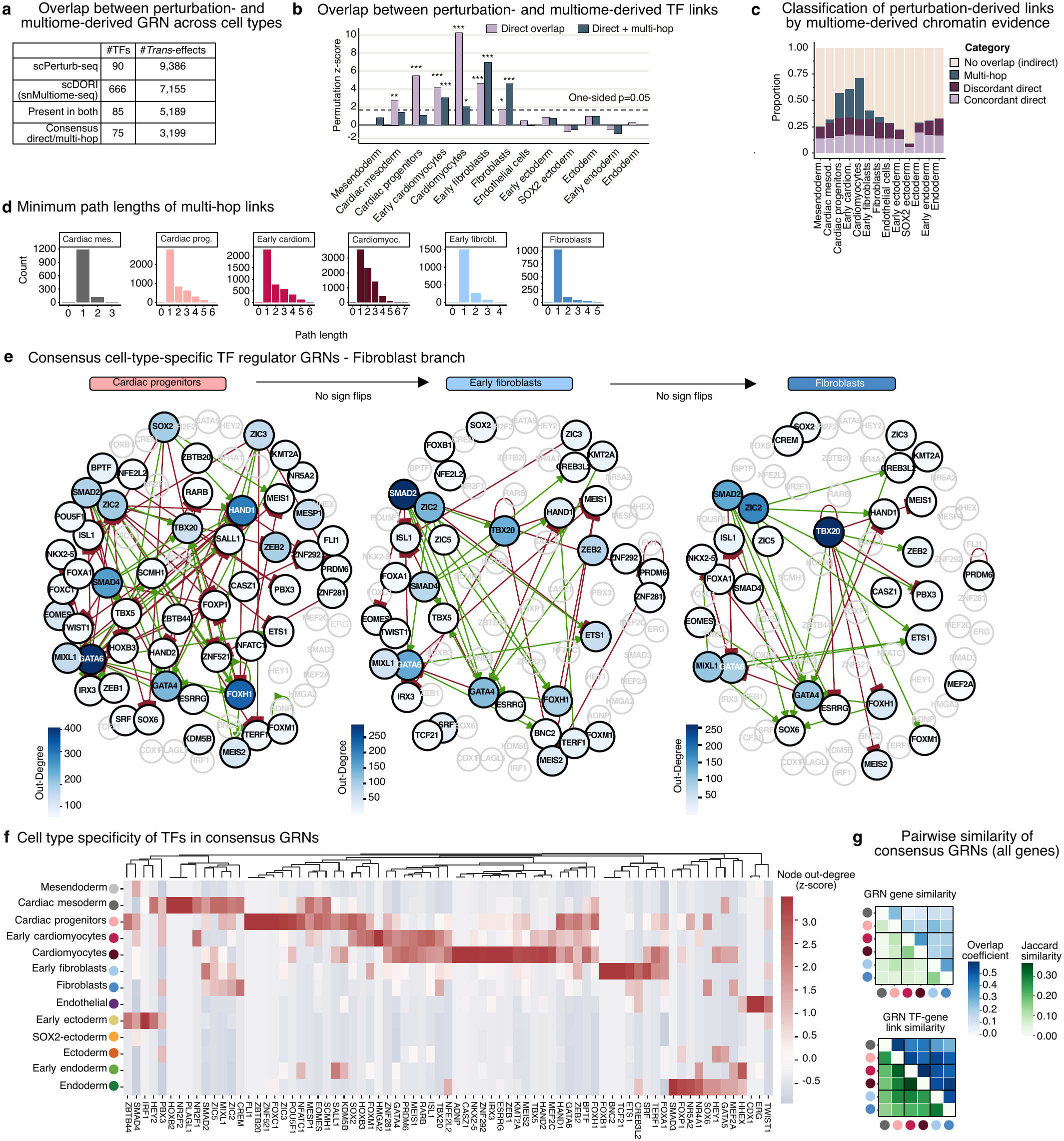
Characterisation of TF-enhancer-gene regulatory network links. a, Number of TFs and *trans*-effects overlapping between scPerturb-seq differential expression testing and multiome-derived eGRNs across all cell types, as well as number of TFs and *trans*-effects in consensus GRNs. **b,** Permutation-based assessment of the statistical significance of the overlap between snMultiome-derived GRNs and scPerturb-seq GRN edges. Null distributions were generated by randomly sampling snMultiome-derived GRNs edges with replacement, without preserving TF out-degree. **c,** Stacked barplot of scPerturb-seq-derived links as proportion for each cell type, classified into sign-discordant/-concordant direct (TF-enhancer-gene link) versus multi-hop (TF-enhancer-TF-enhancer-gene) versus indirect links (significant scPerturb-seq *trans*-effects). **d,** Histograms of minimum path lengths of multi-hop links for each cell type with enough power, indicating the number of intermediate regulatory steps connecting TFs to their downstream genes. **e,** TF-TF consensus GRNs as in Fig. 3c, for the fibroblast branch. Nodes, TFs coloured by out-degree (number of downstream genes); edges, activating (green) or repressing (red) TF-TF links; faded grey, TFs connected only in other cell types; labels, links that switch sign between consecutive cell types (TF-only consensus GRN; 72 TFs with consensus links in the cardiomyocyte or fibroblast branch). **f,** Heatmap showing node out degree (z-scored) for TF regulators across cell types. **g,** Heatmap showing Jaccard index (lower triangle) and overlap coefficient (Szymkiewicz-Simpson coefficient, upper triangle) of nodes (TF and genes, top) and TF-gene links (bottom) between GRNs in cardiac lineage cell types.

**Extended Data Figure 6.**
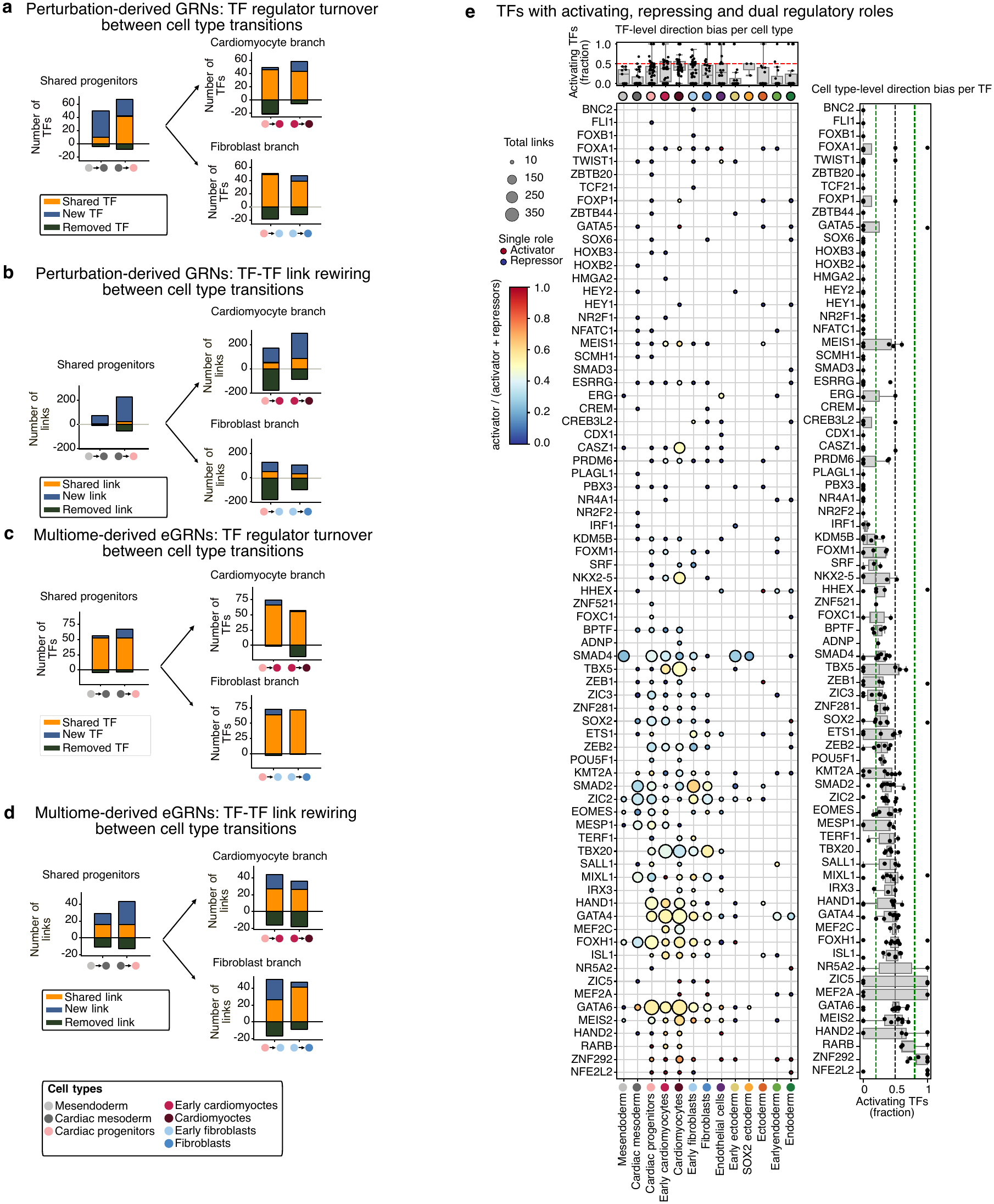
GRN rewiring and activating vs repressing TFs. a-d, TF turnover **(a, c)** and TF-TF link rewiring **(b, d)** between differentiating cell types (left to right) for the cardiomyocyte (top) and fibroblast (bottom) branches. The y-axis shows the total number of TFs regulators **(a,c)** and TF-TF links **(b, d)** for perturbation-derived GRNs (a,b) or multiome-derived enhancer-mediated GRNs (eGRNs) **(c, d)**. Coloured dots on the x-axis mark cell type transitions. Bar colours indicate TFs regulators **(a, c)** or TF-TF links (b,d) that are shared, gained (new) or removed between two consecutive cell types. **e,** TFs with activating, repressing or dual regulatory roles across cell types. Dot plot showing, for each TF (rows) and cell type (columns), the number of regulator links (dot size) and the fraction of activating regulatory links among all links (dot colour) from repressor-dominated (blue, 0), activator–repressor balanced (yellow, 0.5) to activator-dominated (red, 1). Top, boxplots showing the TF-level distribution of the fraction of activating regulatory links within each cell type; dashed line indicates equal activator–repressor balance (ratio = 0.5). Right, boxplots showing the cell type-level distribution of the fraction of activating regulatory links for each TF across all cell types in which it acts; green lines indicate 0.2 and 0.8 respectively; black dashed line indicates equal activator–repressor balance ratio = 0.5.

**Extended Data Figure 7.**
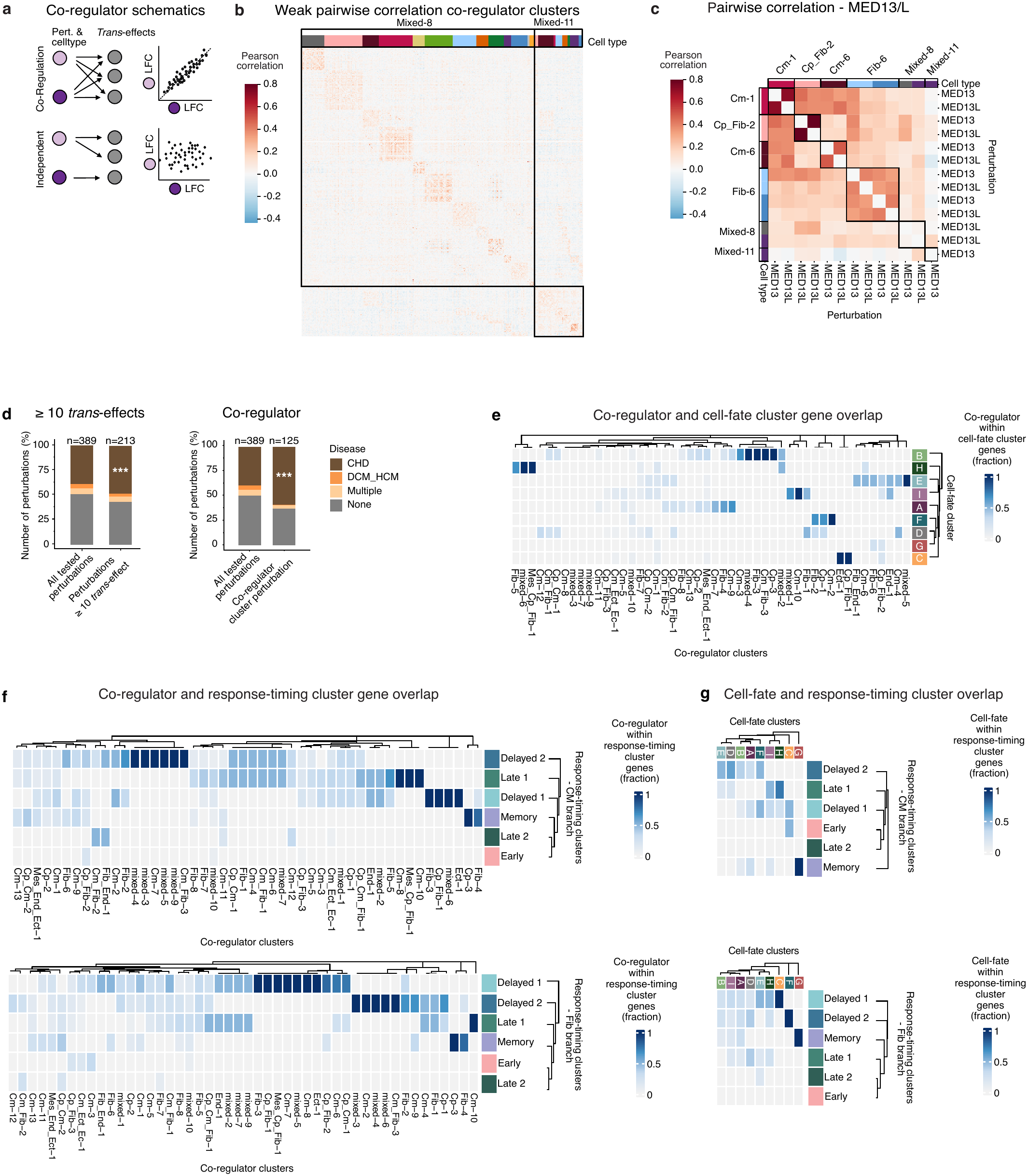
Co-regulator analysis and overlap between cell-fate, co-regulator and response-timing cluster perturbations. a, Schematic of how co-regulators are identified. Co-regulation is defined as correlation of z-score of log-fold changes (LFCs) of *trans*-effects upon perturbation in a cell-type/perturbation combination. **b,** Pairwise Pearson correlation (same as in Fig. 4a) shown for two co-regulator clusters with weak pairwise correlation that were omitted from Fig. 4a. The colour bar on top indicates cell types. Co-regulator clusters are labelled on top. **c,** Zoom-in view highlighting selected regulators, revealing a strong positive correlation between *MED13* and *MED13L*, consistent with their known membership in the same protein complex. Column labels denote co-regulator clusters, colour bars indicate cell types. **d,** Disease annotations for perturbations with ≥ 10 *trans*-effects and perturbations belonging to co-regulators. One-sided Fisher’s exact test for enrichment. ≥ 10 *trans*-effect & CHD \*\*\**P* = 1.50 x 10^-5^, Co-regulator & CHD \*\*\**P* = 5.354 x 10^-8^. n, number of perturbations in each group. **e,** Overlap between cell-fate and co-regulator clusters, showing fraction of co-regulator cluster genes within cell-fate cluster genes. **f, g,** Overlap between co-regulator and response-timing cluster perturbations **(f)** and cell-fate and response-timing cluster perturbations **(g)**, of the cardiomyocyte (top) and fibroblast (bottom) branch. Fractions of overlap of co-regulator/cell-fate cluster perturbations within response-timing perturbations is shown.

**Extended Data Figure 8.**
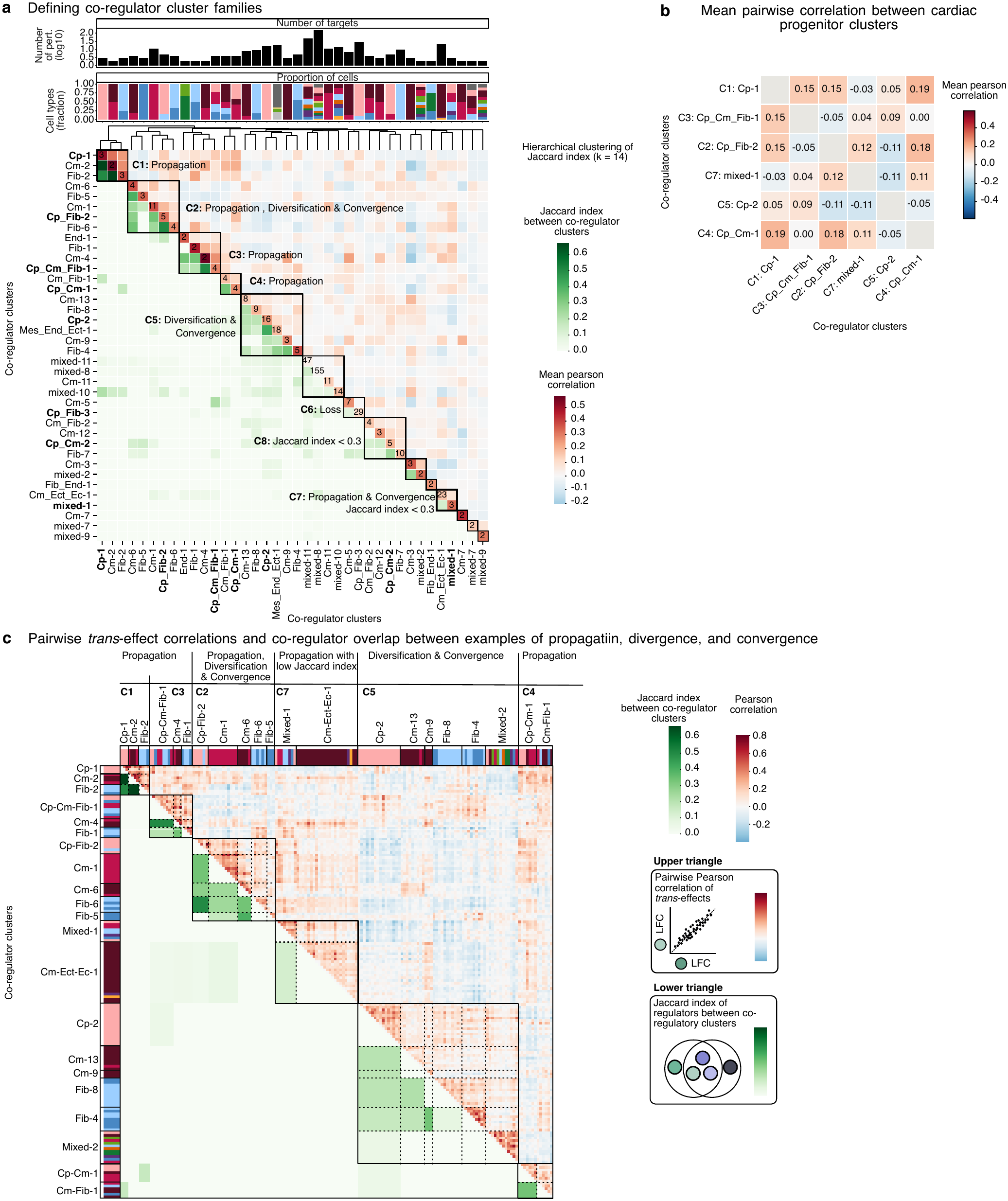
Hierarchical clustering of co-regulator clusters. a, Hierarchical clustering (*k* = 14) of pairwise Jaccard index, between all co-regulator clusters from Fig. 4a, to identify co-regulator cluster families with overlapping regulators. Top bar plot and numbers along diagonal: number of perturbations with *trans*-effects per co-regulator cluster (log10 in bar plot). Bottom stacked bar plot: proportion of cell types comprising each co-regulator cluster. Dendrogram: hierarchical clustering tree used to define the 8 co-regulator families (C1-C8). Families containing only a single co-regulator cluster are excluded from downstream analysis. Heatmap, lower triangle: Jaccard index of shared regulators between co-regulator cluster pairs (green scale); numbers on the diagonal indicate the count of shared regulators within a co-regulator cluster. Heatmap, upper triangle: mean pairwise Pearson correlation of *trans*-effects (z-score of log2 fold-change) between co-regulator clusters (red-blue scale). Co-regulator cluster names in bold along the axes correspond to co-regulators in cardiac progenitors. **b,** Mean pairwise Pearson correlation of *trans*-effects between cardiac-progenitor-containing co-regulator clusters (excluding C6 and C8) highlight examples of broadly similar (positive correlation) or antagonistic (negative correlation) transcriptional programmes affected by specific groups of perturbations. **c,** Heatmap comparing coordinated transcriptional effects across cell types and differentiation states. Upper triangle: pairwise Pearson correlation (red-blue) of *trans*-effects (z-scored log2 fold-change, LFC) between co-regulator clusters. Lower triangle: Jaccard index (green) of shared regulators between co-regulator clusters (diagonal Jaccard index is set to zero). Rows and columns are perturbation-cell type combinations, with labelled black boxes corresponding to co-regulator clusters from Fig. 4a and colour bars indicating cell type. Black boxes drawn on the heatmap denote co-regulator cluster families. Examples show effects that are propagated (C1, C2, C3, C4, C7), diversified (C2, C5) or convergent (C2, C5) during differentiation.

**Extended Data Figure 9.**
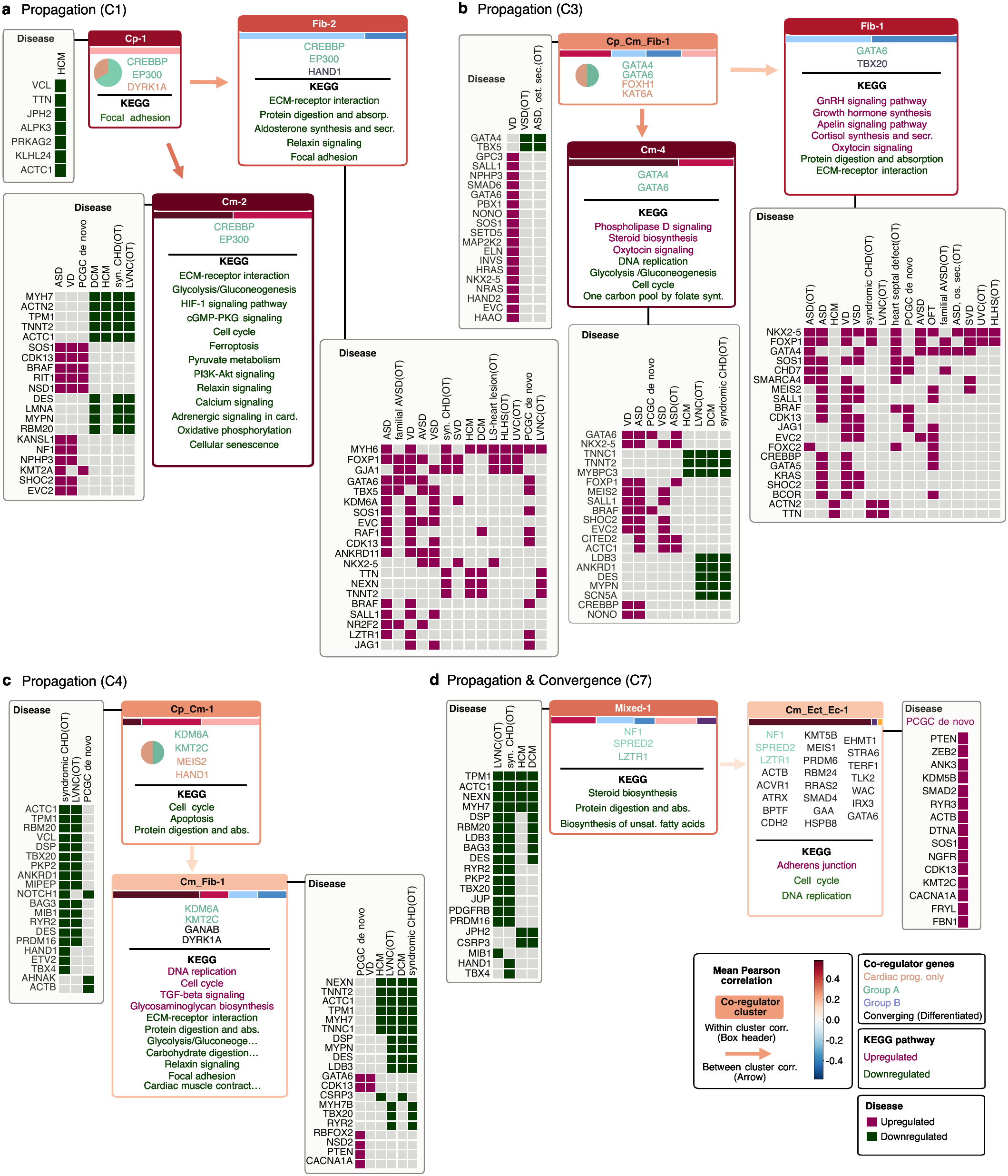
Regulator, pathway and disease-gene annotation of propagation effects. a-d, Plots as in Fig. 5d-f. Propagation in co-regulator family C1 **(a)**. Propagation in co-regulator family C3 **(b)**. Perturbation of *GATA4*, *GATA6*, *FOXH1* and *KAT6A* produces a coordinated transcriptional response in a joint cardiac progenitor/cardiomyocyte/fibroblast state (Cp-Cm-Fib-1) that is conserved after differentiation into both cardiomyocytes (Cm-4, retaining *GATA4* and *GATA6*) and fibroblasts (Fib-1, retaining *GATA6* and additionally coordinating with *TBX20*), each acquiring lineage-specific disease-gene associations and up-or downregulated KEGG pathways. Propagation in co-regulator family C4 **(c)**. Perturbation of *HAND1*, *KMT2C*, *MEIS2* and *KDM6A* produces a coordinated transcriptional response in a joint cardiac progenitor/cardiomyocyte state (Cp-Cm-1) that persists after further differentiation into a joint cardiomyocyte/fibroblast state (Cm-Fib-1), which retains the *KMT2C*/*KDM6A* pairing, loses *HAND1* and *MEIS2*, and gains *DYRK1A* and *GANAB* along with additional KEGG pathways and disease-gene associations. Propagation and convergence in co-regulator family C7 **(d)**.

**Extended Data Figure 10.**
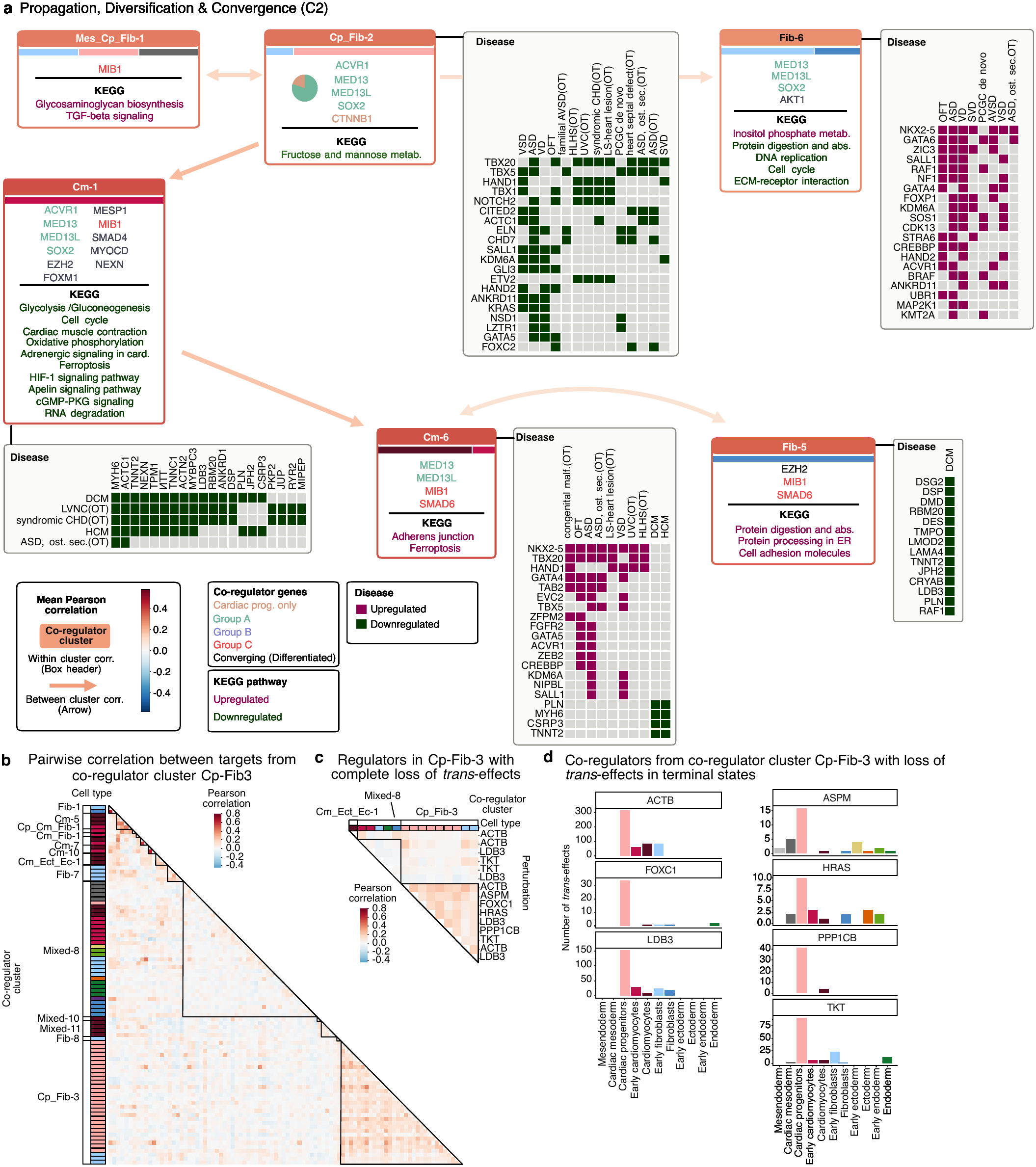
Co-regulator family C2 showing propagation, diversification and convergence effects. a, Plot as in Fig. 5d-f. Propagation, diversification and convergence in co-regulator family C2. Perturbation of ACVR1, CTNNB1, MED13, MED13L and SOX2 produces a coordinated transcriptional response in a joint cardiac progenitor/fibroblast state (Cp-Fib-2). Upon differentiation, this response diversifies along two lineages: In cardiomyocytes it is retained as Cm-1 (keeping ACVR1, MED13, MED13L and SOX2, and gaining EZH2, FOXM1, MIB1, MYOCD, MESP1, NEXN and SMAD4), and in fibroblasts as Fib-6 (keeping MED13, MED13L and SOX2, and gaining AKT1). Continued cardiomyocyte maturation carries Cm-1 into Cm-6 (retaining MED13, MED13L and MIB1, and gaining SMAD6), which then converges with a distinct fibroblast state, Fib-5, sharing MIB1 and SMAD6 (and, more distally, EZH2 with Cm-1) despite Fib-5’s divergent lineage origin. MIB1 is also weakly correlated with group A in Cp-Fib-2. Each cluster gains its own disease-gene associations and up-or downregulated KEGG pathways as indicated. **b,** Pairwise Pearson correlation of trans-effects between co-regulators in co-regulator cluster Cp-Fib-3, including all other co-regulator clusters in which they appear, shown as a lower-triangle heatmap. Row annotations and black outlines drawn on heatmap demarcate co-regulator clusters. **c,** Pairwise correlation of trans-effects show loss of trans-effects in terminal cell states. Top annotation bars indicate co-regulator cluster identity and cell type. **d,** Number of trans-effects across cell types for seven representative co-regulators from co-regulator cluster Cp-Fib-3 (ACTB, ASPM, FOXC1, HRAS, LDB3, PPP1CB, TKT) that show large trans-effects in cardiac progenitor states and near loss of trans-effects in terminal, differentiated cell types.

